# Suppression of transposon mobilization by m^6^A-mediated RNA sequestration in stress granules

**DOI:** 10.1101/2022.03.22.485398

**Authors:** Wenwen Fan, Ling Wang, Zhen Lei, Jie Chu, Jungnam Cho

## Abstract

Transposon is a mobile and ubiquitous DNA that can vastly causes genomic alterations. In plants, it is well documented that transposon mobilization is strongly repressed by DNA methylation; however, the roles of RNA methylation in transposon control remain unknown. Here we suggest that transposon RNA is marked by m^6^A RNA methylation and is sequestered in stress granule (SG) in m^6^A-dependent manner. Intriguingly, a SG-localized AtALKBH9B selectively demethylates a heat-activated retroelement *Onsen*, and thereby releases from spatial confinement allowing for its mobilization. In addition, we show evidence that m^6^A RNA methylation contributes to transpositional suppression by inhibiting the virus-like particles assembly and extrachromosomal DNA production. In summary, this study unveils a hidden role for m^6^A in the suppression of transposon mobility and provides an insight into how transposon counteracts the host’s epitranscriptomic control by hitchhiking RNA demethylase.

## Introduction

Transposable element (TE or transposon) is a DNA that can move from one place to another and is widespread in most eukaryotic genomes (Feschotte, 2008; Lisch, 2012; Rebollo et al., 2012). It consists of two classes; class I DNA transposons that transpose by a ‘cut-and-paste’ mechanism and class II retrotransposons that mobilize by a ‘copy-and-paste’ mechanism through an RNA intermediate (Lisch, 2012; Satheesh et al., 2021). Due to their potentially adverse effects to host genomes caused by transposition, most transposons are strongly repressed by the epigenetic mechanisms including DNA methylation and histone modifications (Law and Jacobsen, 2010; Lisch, 2009; Matzke and Mosher, 2014; Slotkin and Martienssen, 2007; Zhang et al., 2018). Despite the strong epigenetic repression, transposons can be reactivated by environmental challenges, which thereby contribute to genetic diversity and adaptive changes of evolution (Lisch, 2013; Madlung and Comai, 2004; Negi et al., 2016). For example, a Ty1/Copia-like retrotransposon of *Arabidopsis* called *Onsen* can be transcriptionally activated by heat stress and confers heat responsiveness to genes located downstream of insertion positions (Cavrak et al., 2014; Ito et al., 2011). Intriguingly, we in fact observed that the increase of *Onsen* copy number was positively correlated with stronger heat tolerance regardless of its insertion positions, potentially indicating a possible long-distance regulation of heat-responsive genes by *Onsen* in the *Arabidopsis* genome (Fig. S1).

It is well documented that *Onsen* is strongly suppressed by the epigenetic pathways involving small interfering (si) RNAs (Ito et al., 2011; Matsunaga et al., 2012, 2015). Recently, *CHROMOMETHYLASE 3* (*CMT3*) was suggested to promote *Onsen* transcription by preventing CMT2-mediated CHH (H; A, T or C) methylation and histone H3 lysine 9 di-methylation (H3K9me2) accumulation at *Onsen* chromatin under heat stress (Nozawa et al., 2021). Similarly, histone H1 represses the expression of *Onsen* under heat and is required for DNA methylation (Liu et al., 2021b). Whereas the repression of *Onsen* at transcriptional level by DNA methylation is very well characterized, the regulation of *Onsen* RNA at post-transcriptional level has been investigated scantily.

Post-transcriptional RNA modification has emerged as a critical regulatory mark relevant to a variety of RNA processes (Meyer and Jaffrey, 2017; Pan, 2013; Roundtree et al., 2017; Yue et al., 2019). Its study is often referred to as epitranscriptomics analogous to epigenetics. In fact, cellular RNAs contain at least 100 different kinds of post-transcriptional modifications and N6-methyladenosine (m^6^A) is the most abundant modification type present in mRNAs. In plants and other eukaryotes, m^6^A methyltransferases catalyze RNA methylation at a highly conserved motif RRACH (R; G or A) (Luo et al., 2014; Shen et al., 2016). Importantly, several studies have suggested that m^6^A RNA modification regulates TEs; in mammalian cells, for instance, the m^6^A writer complex and reader protein YTHDC1 suppress the expression of endogenous retroviruses (Chelmicki et al., 2021; Liu et al., 2021a). Oppositely, Methyltransferase-like Protein 3 (METTL3) promotes the transposition of Long interspersed element-1 (L1), while RNA demethylase AlkB homolog 5 (ALKBH5) inhibits L1 mobility (Hwang et al., 2021; Xiong et al., 2021).

*Arabidopsis* genome contains 13 ALKBH homologous proteins (Mielecki et al., 2012), of which 5 exhibit high level of similarity to ALKBH5 (Duan et al., 2017). So far only two proteins, AtALKBH9B and AtALKBH10B, have been demonstrated for their catalytic activity of demethylation (Duan et al., 2017; Martínez-Pérez et al., 2017). *AtALKBH10B* is involved in flowering time regulation and directly targets the transcripts of *FT*, *SPL3* and *SPL9* (Duan et al., 2017). *AtALKBH9B* is distinctively expressed in cytoplasm unlike other RNA demethylases (Mielecki et al., 2012). Previous studies have suggested that *AtALKBH9B* negatively regulates the infection of Alfalfa Mosaic Virus (AMV) (Alvarado-Marchena et al., 2021; Martínez-Pérez et al., 2017, 2021). Interestingly, AtALKBH9B co-locates with stress granule (SG) and cytoplasmic siRNA body markers (Martínez-Pérez et al., 2017), potentially implying a functional association with the RNA-mediated epigenetic silencing or RNA degradation pathways. Given the similarity of replication cycles between retrovirus and retrotransposon, TEs might also be subject to the RNA methylation-mediated control; however, transposon regulation by m^6^A RNA modification has been unexplored in plants. In this study, we investigated the role of m^6^A RNA modification in the control of transposon suppression, which involves TE RNA sequestration in SGs.

## Results

### *Onsen* RNA is m^6^A-modified

It is well documented that plant transposons are massively derepressed by heat stress (Cho et al., 2019; Sanchez and Paszkowski, 2014). Despite the strong transcriptional activation of TEs under heat, transposition events are rarely observed (Gaubert et al., 2017; Ito et al., 2011; Thieme et al., 2017), implicating possible repressive mechanisms at RNA level. Since previous studies in human revealed the relevance of m^6^A regulation in the control of retrotransposons, we hypothesized that plant transposons might be controlled in a similar way. In order to test if m^6^A RNA modification plays a role in transposon regulation, we carried out m^6^A-RNA immunoprecipitation sequencing (RIP-seq) experiments using the *Arabidopsis* floral buds harvested before and after 3 hours of heat treatment. Figure 1a shows the distribution of m^6^A mark across transcribed regions, exhibiting a strong peak around the stop codon and a weaker peak at the transcriptional start site, which is consistent with the previously well-known pattern of m^6^A (Arribas-Hernández et al., 2018; Duan et al., 2017; Luo et al., 2014; Shen et al., 2016; Wang et al., 2017). Our m^6^A-RIP-seq data identified almost 2,000 methylated RNAs specific to the heat stress condition (Fig. 1b), which is significantly over-represented with transposons including a retrotransposon family known as *Onsen* (Fig. 1c). *Onsen* is a heat-activated retrotransposon and can produce extrachromosomal (ec) DNA, a pre-integrational reverse-transcribed product of DNA intermediate, and mobilize to new genomic positions (Cho et al., 2019; Ito et al., 2011). *Onsen* exhibited high level of m^6^A along its RNA with the most prominent peak close to the start codon (Fig. 1d). *Arabidopsis* Col-0 genome contains eight intact elements of *Onsen*, and all these elements displayed strong m^6^A enrichment levels (Fig. 1e). We further verified our m^6^A-RIP-seq data by qPCR targeting the upstream regions of *Onsen*, and a strong m^6^A enrichment was detected in regions B and C, which is consistent to Fig. 1d. In summary, the heat-induced transposons including *Onsen* are strongly marked by m^6^A RNA methylation.

**Fig. 1.**
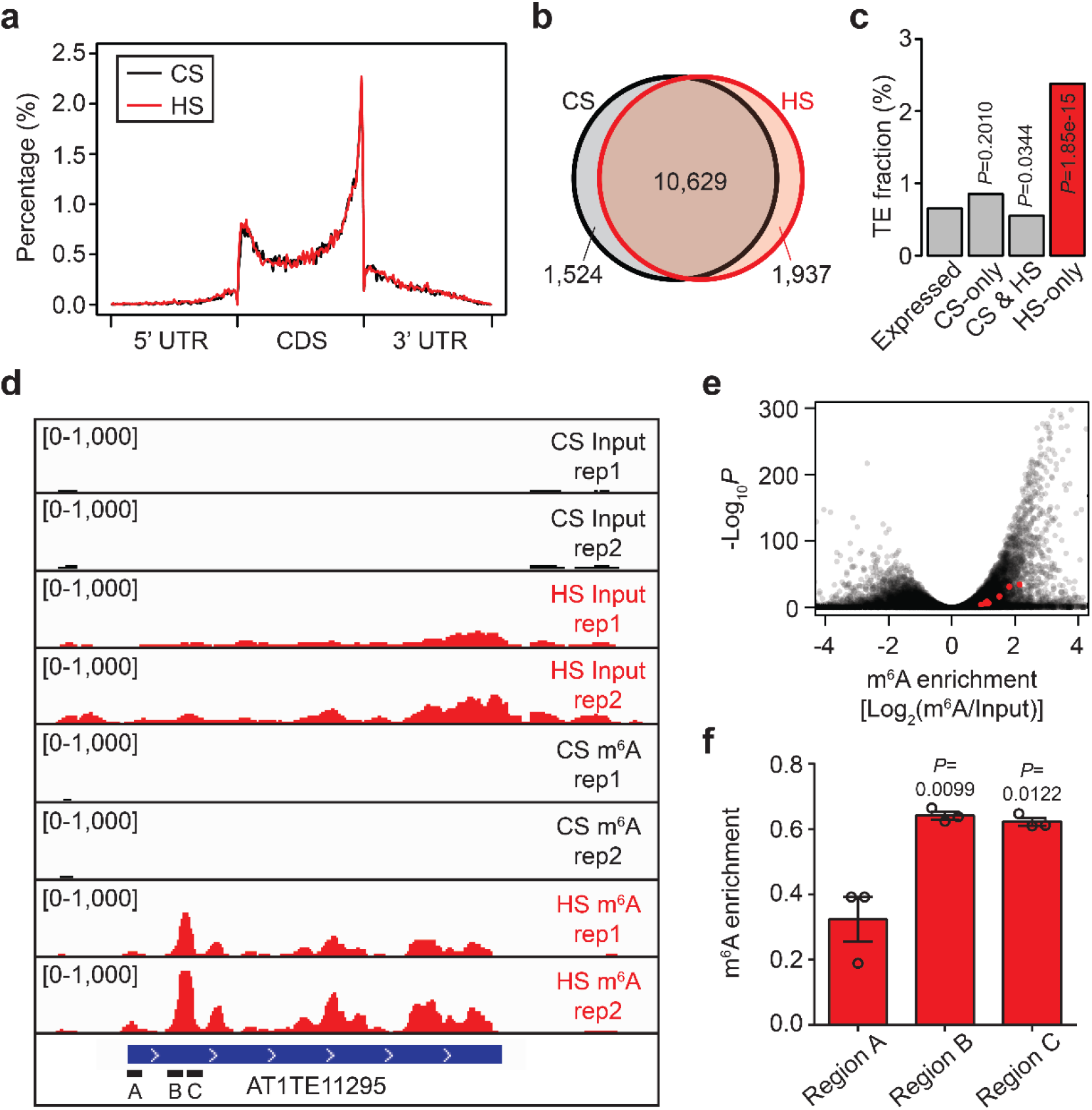
*Onsen* RNA is m^6^A-modified. **a**. Distribution of m^6^A RNA modification in 5’ UTR, CDS, and 3’ UTR. The m^6^A enrichment was calculated for regions spanning 1% of total length. HS, heat-stressed sample; CS, control sample. **b**. Overlap of m^6^A-containing transcripts in CS and HS. m^6^A-modified transcripts were defined as those containing m^6^A peaks detected by MACS2 at FDR lower than 0.05. **c**. The fraction of transposons in each category presented in b. *P* values were obtained by the one-tailed Student’s t-test. **d**. m^6^A-RIP-seq showing *Onsen* locus. **e**. Volcano plot of m^6^A enrichment. Enrichment score was determined by normalizing the m^6^A levels to input levels. The red dots are individual *Onsen* copies. **f**. Validation of m^6^A enrichment by qPCR. The regions tested are as indicated in d. *Act2* was used as an internal control. Values are mean ± s.d. from three biological replications. *P* values were obtained by the one-tailed Student’s t-test.

### AtALKBH9B is an m^6^A RNA demethylase targeting *Onsen*

In search for possible regulators of m^6^A of *Onsen* RNA, we analyzed the RNA-seq data of Gaubert et al. to profile the expression pattern of m^6^A regulators (Gaubert et al., 2017). Most of the genes encoding for m^6^A writers, erasers, and readers were upregulated upon heat stress (Fig. S2a-c), possibly indicating a functional association of RNA methylation and heat stress response. Among these genes, we paid attention to *AtALKBH9B* (hereinafter *9B*) which was previously suggested to regulate viral RNAs (Martínez-Pérez et al., 2017). Since TE RNAs share several cellular characteristics with viral RNAs, we hypothesized that *9B* might also regulate *Onsen* RNA. In addition, the expression pattern of *9B* and *Onsen* during a time course of heat treatment was similar, displaying a rather slow increase and peak at 24 h after heat stress (Fig. S2d and e).

To test if *9B* is involved in *Onsen* RNA regulation, we first isolated a T-DNA insertional mutant *9b-1* and also generated a deletion mutant *9b-2* using the CRISPR-Cas9 system (Fig. 2a and Fig. S3). The RNA levels of *Onsen* were strongly upregulated in both mutants (Fig. 2b), and similar pattern was also observed in RNA-seq data (Fig. 2c). We then expressed the GFP-tagged 9B in the *9b-1* mutant and could detect a suppression of *Onsen* RNA to the wild-type (wt) level (Fig. 2d). In addition, previous studies suggested that *9B* and *10B* are expressed at high level throughout various developmental stages, while other RNA demethylases are marginally expressed (Duan et al., 2017). The *10b* mutants were thus tested for *Onsen* RNA levels; however, we were not able to detect any noticeable changes, suggesting that *10B* plays a negligible role in *Onsen* RNA regulation (Fig. S4). We next performed m^6^A-RIP-qPCR experiments using the wt and *9b-1* mutant. As shown in Fig. 2e, the m^6^A levels were significantly elevated in the *9b-1* mutant, indicating that 9B might cause demethylation of *Onsen* RNA. To determine if 9B is a direct regulator of *Onsen* RNA, RIP-qPCR experiments were performed using the *p9B::9B:GFP* transgenic plants used in Fig. 2d. Figure 2f shows a substantial level of 9B:GFP enrichment to *Onsen* RNA. These altogether suggest that 9B is an RNA demethylase that directly targets *Onsen* RNA.

**Fig. 2.**
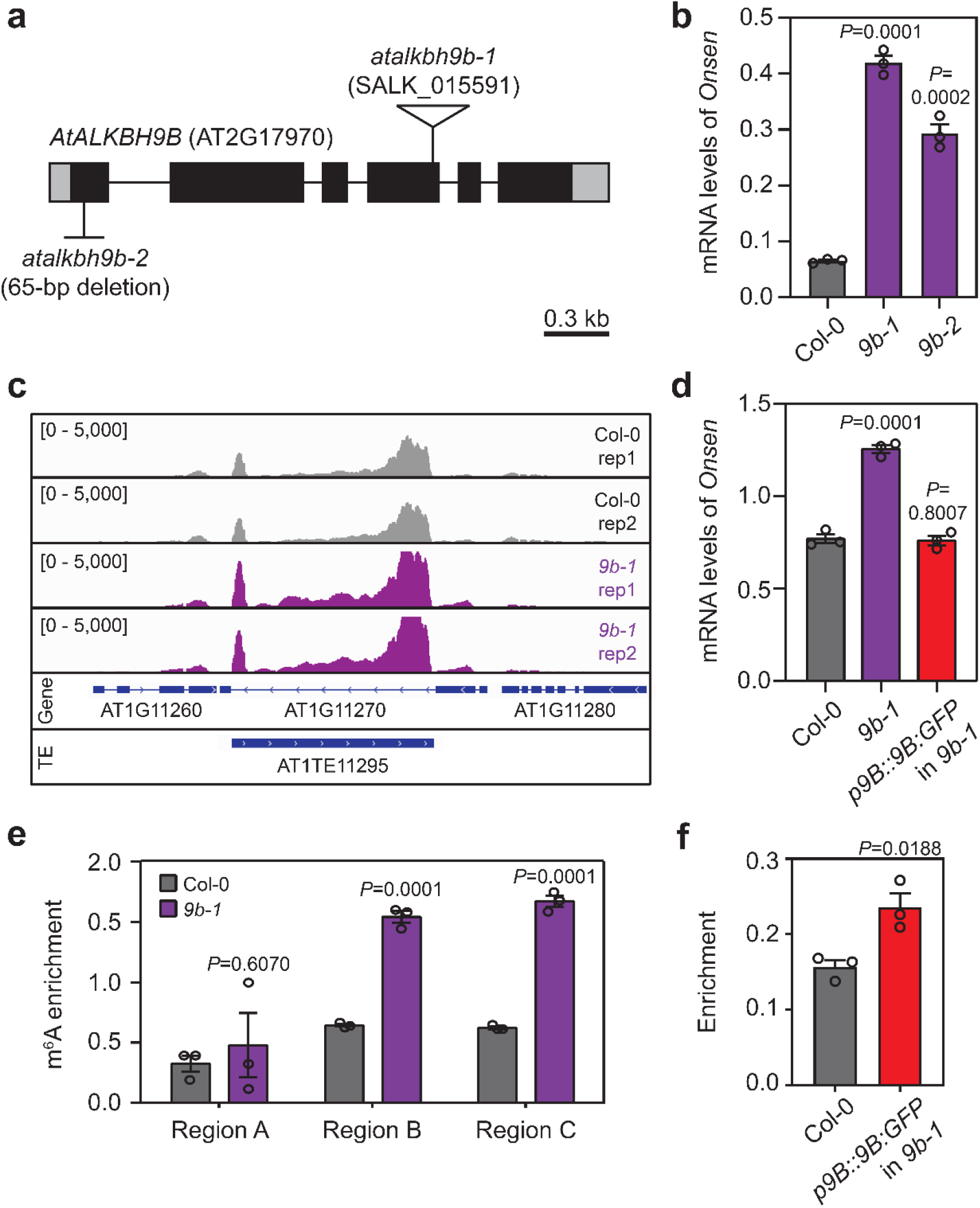
*AtALKBH9B* regulates *Onsen* RNA. **a**. Gene structure of *AtALKBH9B*. T-DNA insertion of *9b-1* is shown as a triangle. The *9b-2* mutant contains a large deletion of 65 bp in the first exon. **b**. *Onsen* RNA levels in the *9b* mutants determined by RT-qPCR. *Act2* was used as an internal control. Values are mean ± s.d. from three biological replications. *P* values were obtained by the one-tailed Student’s t-test. **c**. RNA-seq of *9b-1* showing the *Onsen* locus. **d**. RT-qPCR for complementation assay of the *9b-1* mutant with the *pro9B::9B:GFP* construct. *Act2* was used as an internal control. Values are mean ± s.d. from three biological replications. *P* values were obtained by the one-tailed Student’s t-test. **e**. m^6^A-RIP-qPCR performed in the *9b-1* mutant. Regions are as indicated in Fig. 1d. *Act2* was used as an internal control. Values are mean ± s.d. from three biological replications. *P* values were obtained by the one-tailed Student’s t-test. **f**. RIP-qPCR experiments using the *pro9B::9B:GFP* transgenic plants used in d. *Act2* was used as an internal control. Values are mean ± s.d. from three biological replications. *P* values were obtained by the one-tailed Student’s t-test.

### Reduced transposition of *Onsen* in *9b*

Since the RNA levels of *Onsen* were increased in the *9b* mutants, we speculated that the transpositional activity would be also increased. To directly determine the *Onsen* mobility, we first carried out ALE-seq experiments and assessed the pre-integrational DNA intermediate levels of *Onsen*. Intriguingly, the ecDNA levels of *Onsen* were drastically reduced in the *9b-1* mutant (Fig. 3a). Similar results were also observed in the experiments detecting the total DNA levels of *Onsen* in the heat-stressed *9b-1* mutant (Fig. 3b). We then wanted to directly measure the insertional activity of *Onsen* in the *9b-1* mutant and for this we performed droplet digital PCR (ddPCR) experiments to quantitatively determine the copy numbers (Fan and Cho, 2021). In this experiment, plants were treated with heat stress and drugs (α-amanitin and zebularine), which were known to induce *Onsen* retrotransposition even in the wt background (Thieme et al., 2017). Surviving plants after heat stress were grown to maturity and seeds were separately collected from individual plant. DNA extracted from the next generation plants was subjected to ddPCR that simultaneously amplified *Onsen* and *CBF2*, which was used as a single-copy reference gene. Consistently, the *9b-1* mutant generated fewer number of new *Onsen* copies than wt (Fig. 3c). These data indicate that *9B* is required for *Onsen* retrotransposition.

**Fig. 3.**
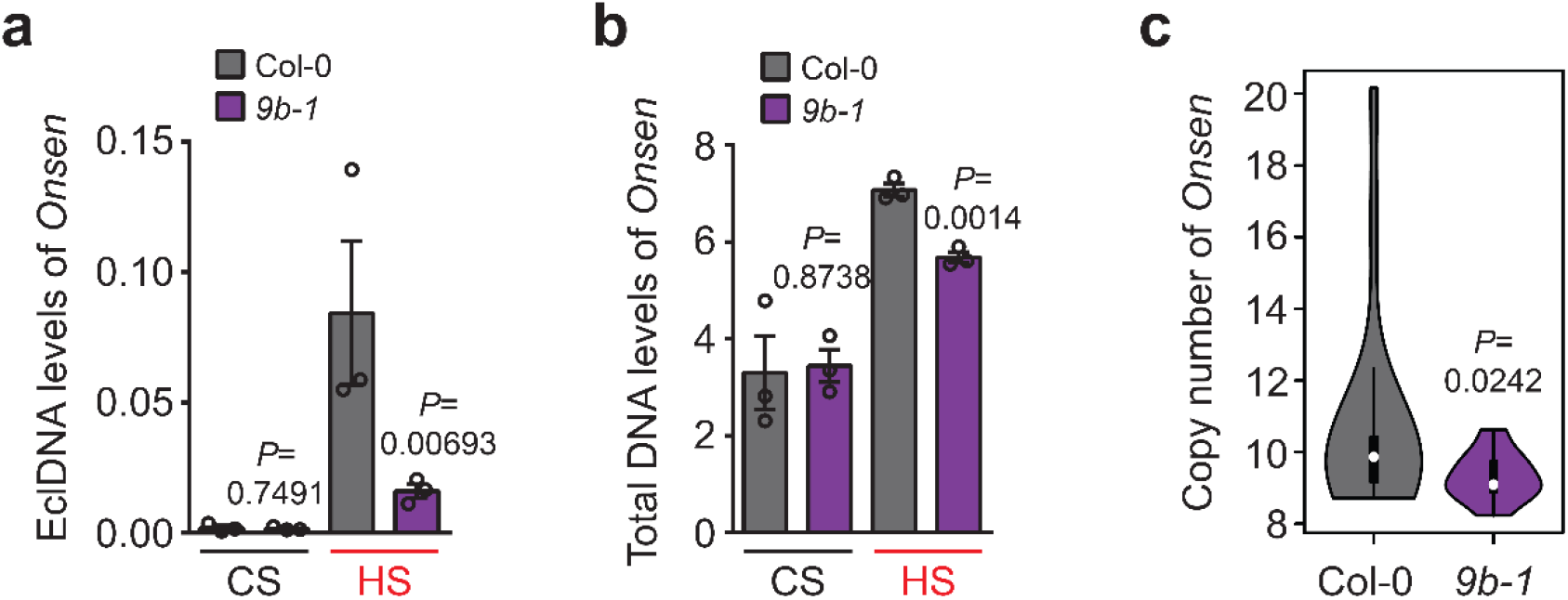
Retrotransposition activity of *Onsen* is reduced in *9b-1*. **a**. The levels of eclDNA of *Onsen* in the *9b-1* mutant. ALE-seq was performed to determine the eclDNA levels. PCR-amplified *Evade* DNA was used as spike-in control. CS, control sample; HS, heat-stressed sample. Values are mean ± s.d. from three biological replications. *P* values were obtained by the one-tailed Student’s t-test. **b**. Total DNA levels of *Onsen* in the *9b-1* mutant. *Act2* was used as an internal control. Values are mean ± s.d. from three biological replications. *P* values were obtained by the one-tailed Student’s t-test. **c**. Copy number of *Onsen* in the progenies of wt (n=20) and *9b-1* mutants (n=20) that were subjected to heat stress treatment. Copy number was determined by ddPCR using *CCR2* as a single-copy control. *P* value was obtained by the one-sided Mann-Whitney U test.

### 9B and m^6^A-methylated RNAs are localized to SGs in heat stress

Our results so far indicated that the *9b* mutant exhibits opposing patterns to different *Onsen* intermediates, i.e., increased RNA and reduced DNA level. We speculated that such divergence was caused probably by RNA sequestration that inhibits the conversion of RNA to DNA intermediates. Previous studies in human demonstrated a strong association of methylated RNAs with SGs (Fu and Zhuang, 2020; Ries et al., 2019). SG is an evolutionary conserved intracellular compartment that is formed under stress conditions and stores proteins and RNAs (Chantarachot and Bailey-Serres, 2018; Kim et al., 2021; Riback et al., 2017; Wheeler et al., 2017). We therefore hypothesized that hypermethylated *Onsen* RNA in *9b* might be sequestered in SG and was precluded from where the ecDNA production happens. To test this hypothesis, we first investigated if 9B protein is localized in SGs. We expressed 9B:GFP in tobacco leaves along with SGS3:TdTomato as a SG marker. As shown in Fig. 4a, cytoplasmic foci of SGs were formed under heat stress, and 9B colocalized well together with SGS3. The association of 9B to SGs was further examined with the double transgenic *Arabidopsis* plants expressing both 9B:GFP and SGS3:TdTomato. Consistent to Fig. 4a, 9B:GFP located to cytoplasmic foci along with SGS3:TdTomato (Fig. 4b and c). We further investigated the interactome of 9B by performing the IP-mass spectrometry (MS) experiments. Our IP-MS data identified many proteins that were previously known as SG components (Fig. 4d, Fig. S5 and Table S3). Importantly, YTH domain-containing m^6^A reader proteins, ECT1 and ECT2, were also identified in the 9B interactome (Fig. 4d, Table S3). It is worth noting that a substantial fraction of 9B-interacting proteins is commonly found in the SG proteome which was reported previously (Table S4) (Kosmacz et al., 2019). Among those proteins that are in both 9B interactome and stress granule proteome, ECT2 was chosen for further test of interaction with 9B. We performed split luciferase assay experiments in tobacco leaves and observed significant interaction between 9B and ECT2 (Fig. 4e). Overall, 9B is localized to SGs and interacts with multiple SG components.

**Fig. 4.**
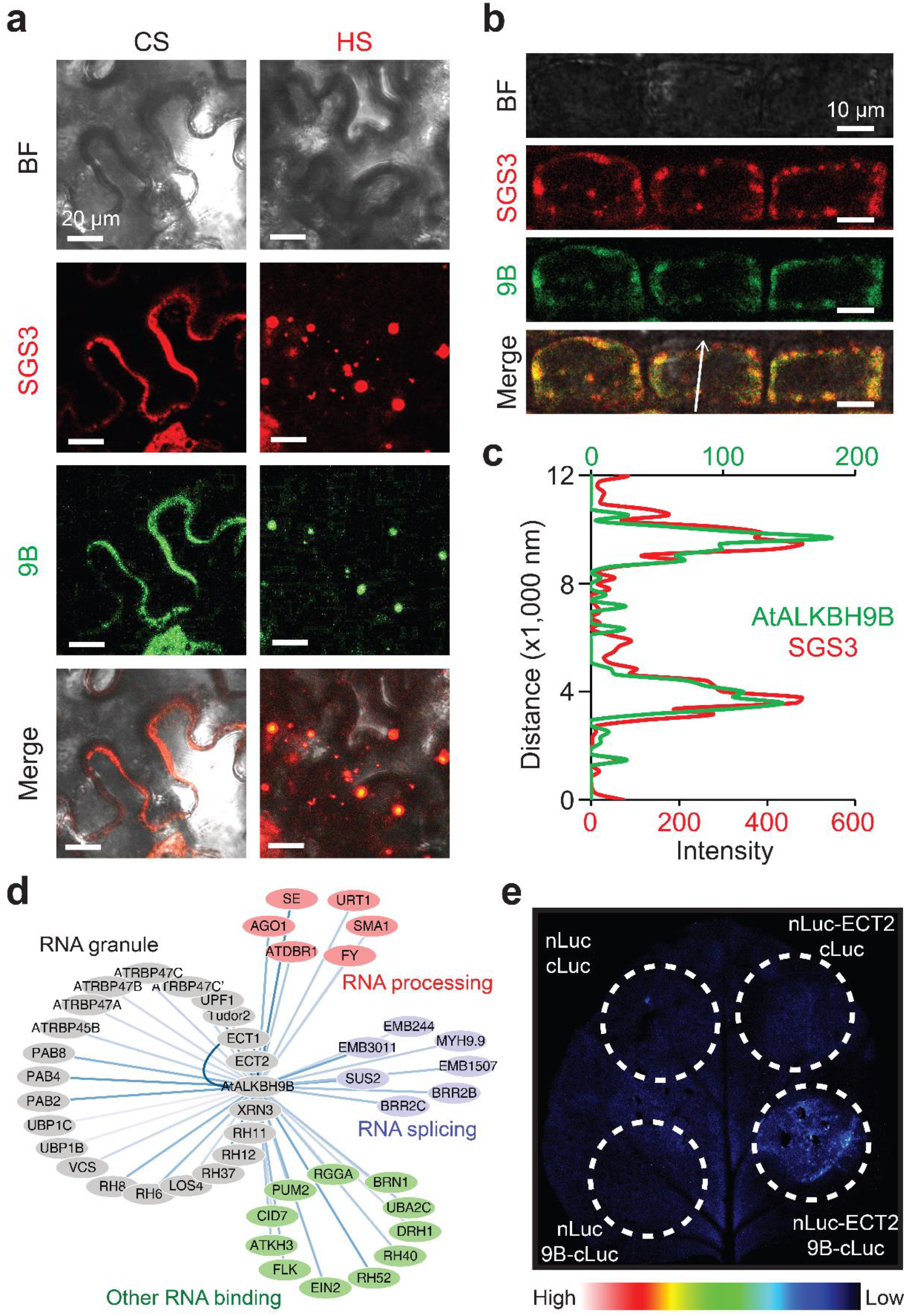
AtALKBH9B is localized to stress granules. **a**. Co-localization of 9B-GFP and SGS3-TdTomato tested in tobacco transient expression system. CS, control sample; HS, heat-stressed for 12 h. Bar=20 µm. **b** and **c**. Co-localization of 9B-GFP and SGS3-TdTomato in *Arabidopsis* double transgenic plants (**b**). Root epidermal cells of heat-stressed plants are shown. Arrow indicates the section for which the signal intensity was quantitated (**c**). Bar=10 µm. **d**. Interactome of 9B protein revealed by IP/MS. The 9B-GFP transgenic *Arabidopsis* plants were used. Proteins are color-coded by their functional categories. Edges with darker color represent stronger protein-protein interactions. **e**. Interaction of 9B and ECT2 determined by a split luciferase assay performed in tobacco transient expression system.

We next investigated if m^6^A-methylated RNAs are preferably localized to stress granules in *Arabidopsis*. For this, we carried out dot blot experiments with m^6^A antibody using the RNAs extracted from the SG fraction of the heat-stressed *9b-1* mutants. Compared to total RNA, SG RNA showed higher level of m^6^A RNA modification, which is consistent to mammalian SG (Fig. 5a and b). RNA-seq was also performed using the total and SG RNAs derived from the heat-stressed wt and *9b-1* mutant. We first identified transcripts upregulated in *9b-1* and determined their SG enrichment. When compared with randomly selected RNAs, the *9B*-regulated transcripts exhibited stronger SG enrichment (Fig. 5c), indicating that *9B* plays a role in the m^6^A-dependent RNA localization to SGs. The enhancement of SG localization of *Onsen* RNA in *9b* was further validated by qPCR. Figure 5d shows that the stress granule enrichment of *Onsen* RNA was increased in the *9b-1* mutant (Fig. 5d). We also tested the mutants for SG components and observed that the *Onsen* RNA levels are decreased in these mutants (Fig. S7), which partly indicates that SG stabilizes *Onsen* RNA. In short, 9B and m^6^A-methylated transcripts are localized in cytoplasmic SGs under heat stress.

**Fig. 5.**
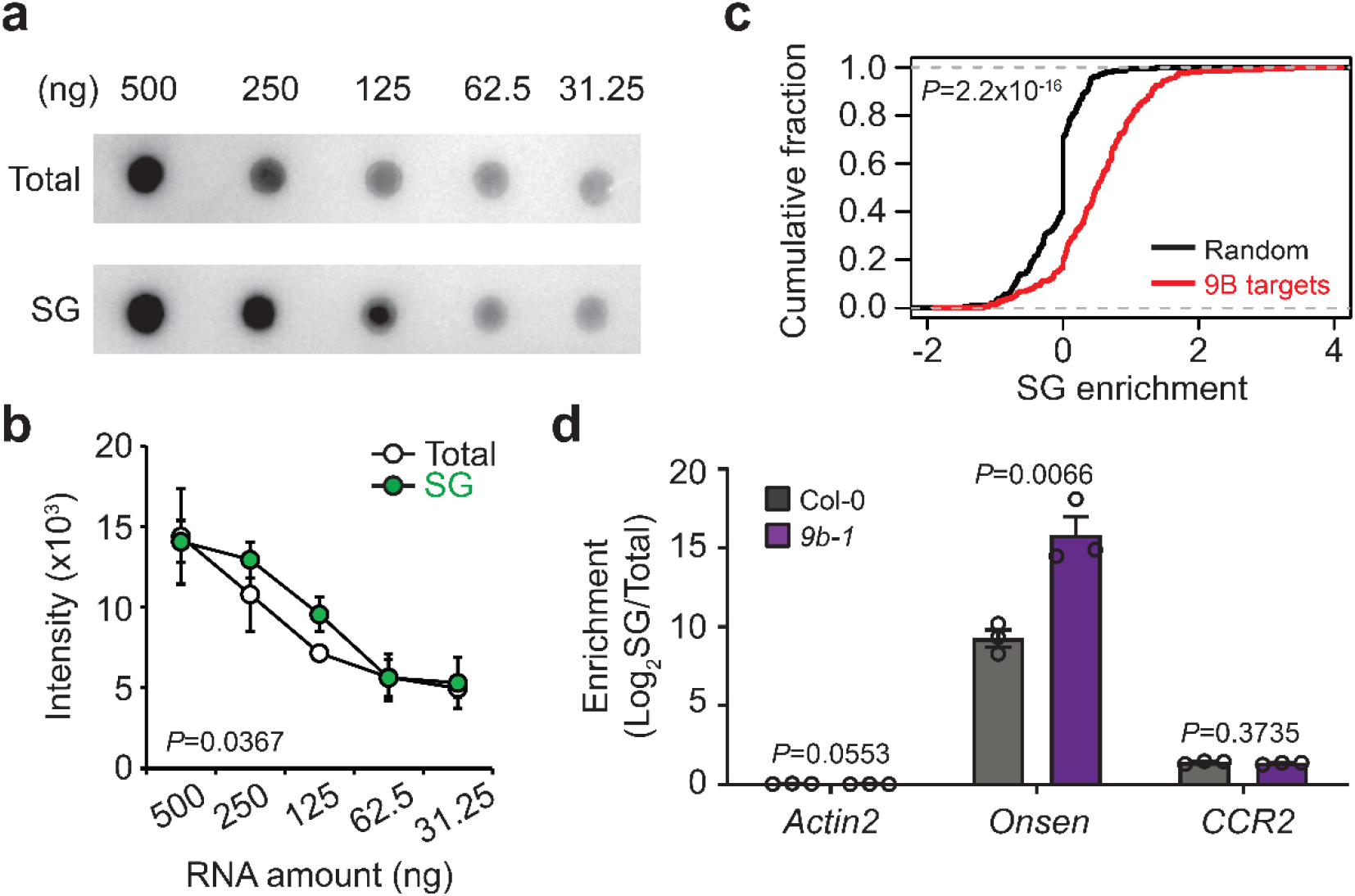
Strong enrichment of m^6^A-modified RNA in SGs. **a**. Dot blot of m^6^A-modified RNA. Total and SG fraction RNA derived from the *9b-1* mutant plants were used. Serially diluted RNA concentration is indicated. **b**. Signal intensity of the blot is quantitated. Values are mean ± s.d. from three biological replications. **c**. Cumulative distribution of the SG enrichment of the 9B target genes as compared with the randomly selected genes. *P* values were obtained by the one-sided Wilcoxon rank sum text. **d**. SG enrichment of *Onsen* RNA in the *9b-1* mutant determined by qPCR. Normalization was against total RNA. *CCR2* was used as a negative control (Fig. S6). Values are mean ± s.d. from three biological replications. *P* values were obtained by the one-tailed Student’s t-test.

### m^6^A inhibits VLP assembly and reverse transcription of ecDNA

We so far demonstrated that m^6^A-methylated *Onsen* RNA is sequestered in SG and 9B demethylates it, allowing for progression to mobilization cycle. It is, however, important to note that neo-insertions of *Onsen* are hardly detectable in the wt background (Ito et al., 2011; Thieme et al., 2017). We thus postulated additional inhibitory effects of m^6^A RNA modification on retrotransposition of *Onsen* other than RNA sequestration to SGs. Since the production of pre-integrational DNA intermediates happens in virus-like particles (VLPs) and the physical interaction between template RNA and retroelement-encoded Gag protein is the first step of it (Cho et al., 2019; Satheesh et al., 2021), we tested if the methylation status of RNA influences the affinity of Gag with RNA. Fluorescence polarization assay was carried out to determine the in vitro binding activity of Gag using the RNA oligonucleotides of identical sequence but differing in m^6^A methylation. As shown in Fig. 6a, m^6^A-modified RNA exhibited the increased *Kd* value compared to non-methylated RNA, suggesting that m^6^A inhibits the binding to Gag. We further examined the interaction of Gag and *Onsen* RNA by performing RIP-qPCR experiments using the Gag-GFP-expressing transgenic *Arabidopsis* plants (Fig. S8). Gag-GFP showed strong binding enrichment to *Onsen* RNA; however, when Gag-GFP was expressed in the *9b-1* mutant background, the binding enrichment was drastically reduced (Fig. 6b). In addition, we also speculated that m^6^A RNA methylation might interfere with reverse transcription process. To test this possibility, partial *Onsen* RNA was in vitro transcribed in presence or absence of m^6^A and subjected to RT-qPCR. Consistent to previous studies suggesting that reverse transcriptional activity can be compromised by RNA modifications (Ryvkin et al., 2013; Woodson et al., 1993), m^6^A-modified RNA showed higher Cq values (Fig. S9) which indicates that RNA methylation inhibited cDNA production. These data altogether suggest that m^6^A RNA methylation contributes to retrotransposition suppression by inhibiting VLP assembly and reverse transcription to ecDNA.

**Fig. 6.**
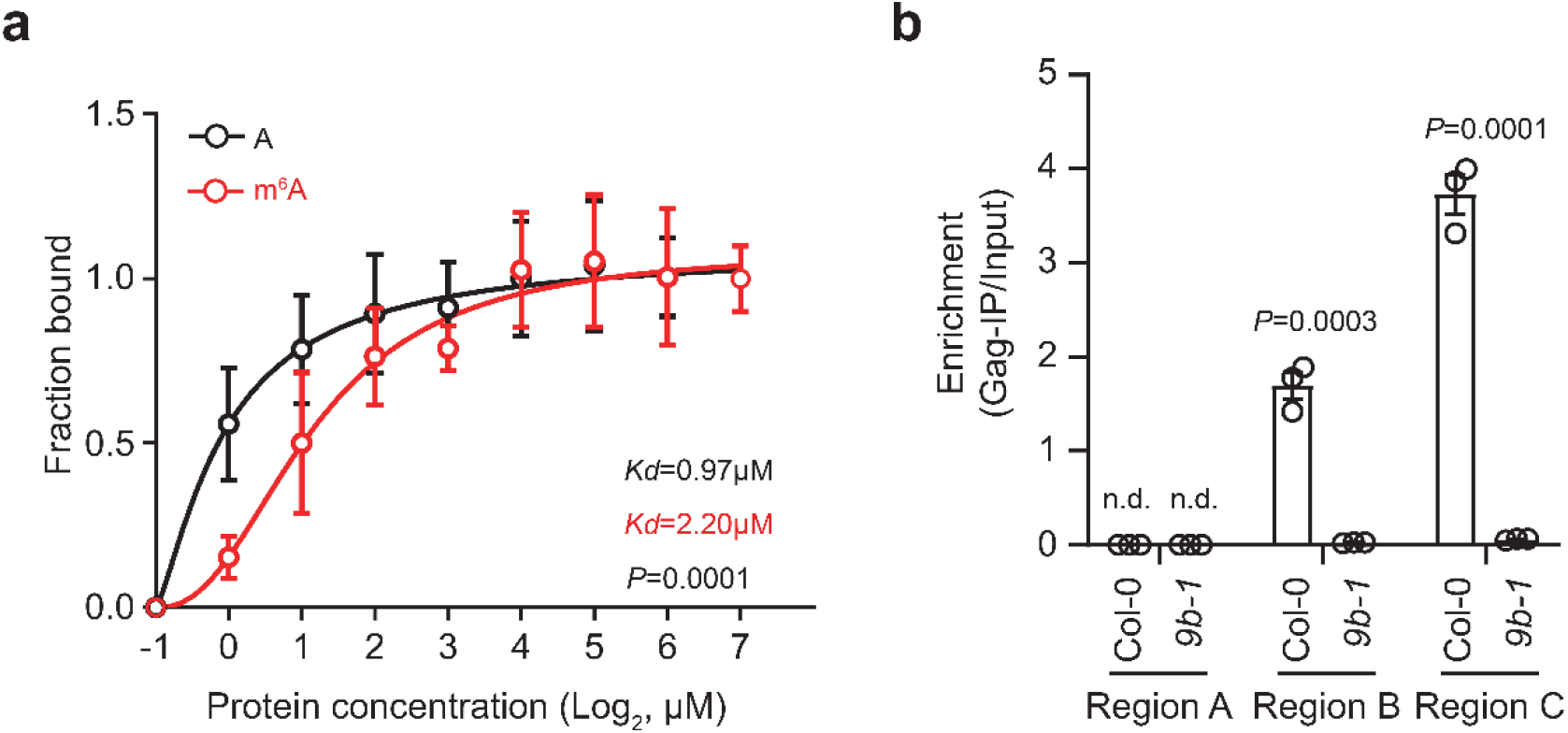
m^6^A RNA methylation inhibits binding of Gag in vitro and in vivo. **a**. Fluorescence polarization assay of purified Gag protein of *Onsen*. 12-mer RNA oligos with or without m^6^A modification (GGCCAACUACGU and GGCCAm^6^ACUACGU) were used. Values are mean ± s.d. from four technical replications. *P* values were obtained by the two-way ANOVA. **b**. RIP-qPCR of Gag-GFP binding to *Onsen* transcript. *OnsenLTR::Gag:GFP* construct was introgressed to *9b-1* by genetic cross. Regions tested are as described in Fig. 1d. Values are mean ± s.d. from three biological replications. *P* values were obtained by the one-tailed Student’s t-test.

## Discussion

In this study we suggested that a heat-activated retrotransposon *Onsen* is m^6^A-methylated at RNA and localized to cytoplasmic SGs (Fig. 7). m^6^A RNA methylation not only leads to spatial constraints preventing RNA maturation to ecDNA, but also biochemically inhibits VLP assembly and reverse transcription process (Fig. 7). Importantly, a SG-localized RNA demethylase AtALKBH9B directly targets *Onsen* RNA and allows it to complete the mobilization cycle. Our study provides an insight into the biological role of SG as a site for seclusion of transposon RNAs and thus suppression of their mobility. This notion is partially in agreement with previous work in mammalians that suggested the m^6^A-mediated inhibition of transposons through RNA destabilization (Chelmicki et al., 2021; Liu et al., 2021a). However, the discrepancy of the roles of m^6^A in *Arabidopsis* from that of animals is that it does not trigger strong RNA decay but is associated with RNA stabilization (Anderson et al., 2018; Luo et al., 2014). Our data also showed that the depletion of SG components results in the reduction of *Onsen* RNA levels (Fig. S7), indicating that SG enhances RNA stability. These altogether indicate that m^6^A RNA methylation of *Onsen* sequesters RNA to SG without compromising its RNA stability.

**Fig. 7.**
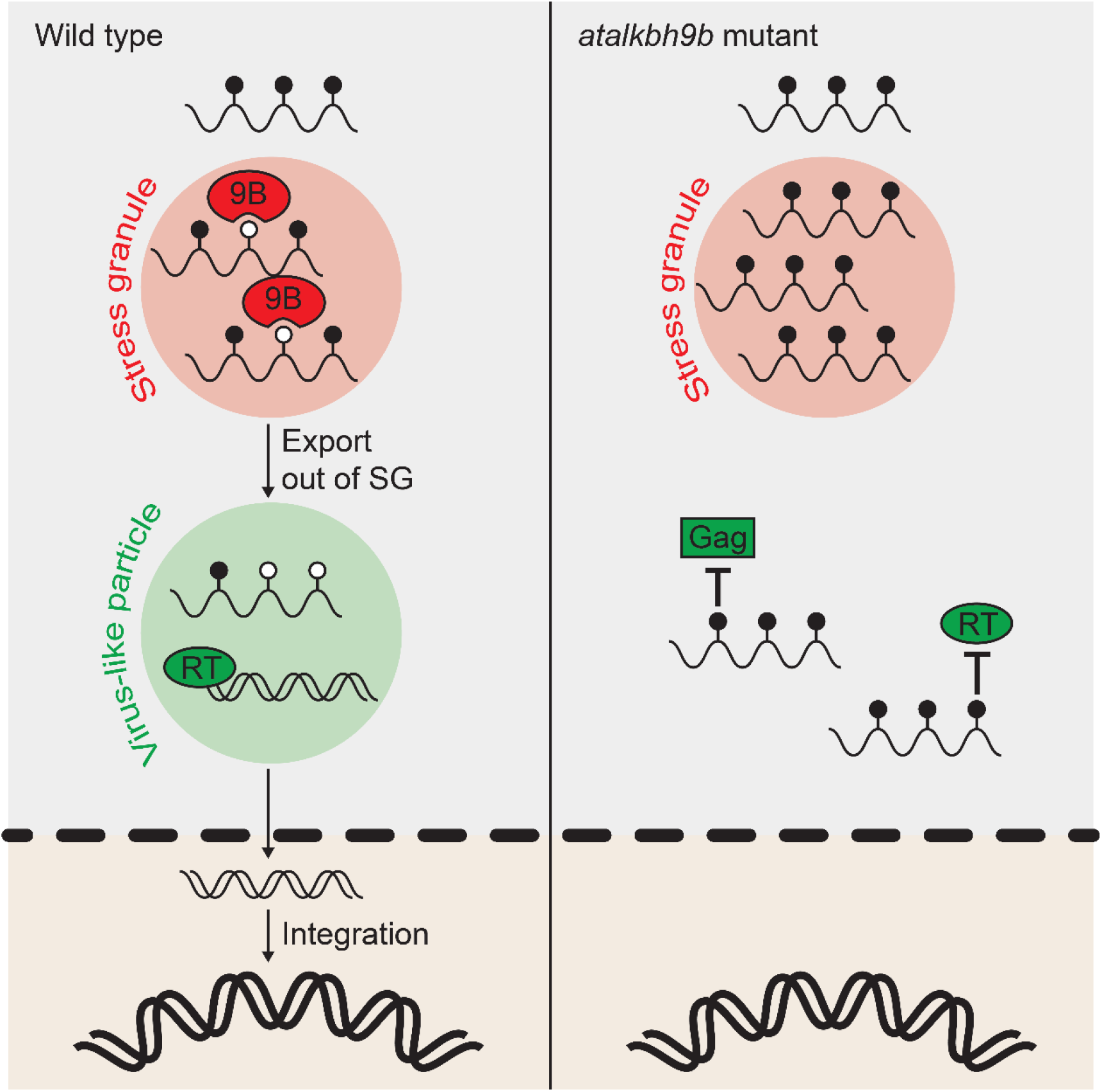
A proposed model. Methylated *Onsen* RNA is localized to SG. AtALKBH9B demethylates and exports out *Onsen* RNA. Hypomethylated *Onsen* RNA assembles to VLP by interacting with Gag protein and is reverse-transcribed to form ecDNA. In the mutant of *AtALKBH9B*, *Onsen* RNA is hypermethylated at m^6^A and sequestered in SG. In addition, RNA methylation inhibits the binding of Gag and reverse transcription.

AtALKBH9B was previously characterized to locate to SGs and facilitate the viral infectivity (Martínez-Pérez et al., 2017), which is similar to what we observed for retrotransposons. In fact, AtALKBH9B is peculiar and distinct from AtALKBH10B, the other active RNA demethylase in *Arabidopsis*. For example, AtALKBH9B is localized to cytoplasmic foci and regulates relatively fewer number of transcripts (Duan et al., 2017; Mielecki et al., 2012). Importantly, we showed that the *Onsen* transcript levels are not altered in the *10b* mutants (Fig. S4). These imply the functional diversification of RNA demethylases and indicate that *AtALKBH9B* has evolved to preferentially target non-native and invasive genetic elements such as retroviruses and retrotransposons. Further investigation to biochemical characteristics of AtALKBH9B will be required to understand how RNA demethylases discriminate their RNA targets. A recent study on the targeting rules of ECT2 can hint us at possible RNA recognition mechanisms of m^6^A regulation (Arribas-Hernández et al., 2021). The ECT2-binding transcriptome revealed a strong sequence bias towards U that is enriched around m^6^A-modified sites (Arribas-Hernández et al., 2021). Interestingly, similar sequence bias of low GC contents in transposons was suggested in our previous report (Kim et al., 2021). Since ECT2 was also identified as one of the interacting proteins of AtALKBH9B in this study (Fig. 4d and e), it will be an interesting follow-up study to investigate if ECT2 has a role in the specific RNA target recognition of AtALKBH9B.

In summary, mobile genetic elements are subject to multi-layered repression at transcriptional and post-transcriptional steps. Our work suggests a previously unknown mechanism to suppress transposon mobility that involves m^6^A RNA methylation and sequestration of TE RNA in SG. In addition, retrotransposon *Onsen* provides an intriguing example of adopting a unique RNA demethylase to bypass such suppression.

## Materials and Methods

### Plant materials and growth condition

*Arabidopsis* ecotype Col-0 was used as wt to compare with mutants including *atalkbh9b-1* (SALK_015591C), *g3bpl-1* (SALK_011708), *sgs3-14* (SALK_001394), and *ago7-1* (SALK_037458). To induce *de novo* mutations in *AtALKBH9B*, a CRISPR-Cas9 vector was constructed by cloning three sgRNAs (sequences are provided in Supplementary Table 1) into the Cas9-containing binary vector. The oligonucleotides encoding the sgRNAs were first cloned into the pENTR_L4_R1, pENTR_L1_L2 and pENTR_R2_L3 entry vectors at the BbsI sites. The entry vectors containing the sgRNAs were then cloned into the destination vector by the LR recombination reaction using the MultiSite Gateway Pro kit (Thermo Fisher Scientific). The resulting vector was transferred to *Agrobacterium tumefaciens* strain GV3101 and transformed into Col-0. As the vector carries a GFP fluorescence gene driven by a seed coat-specific promoter, the collected T1 seeds were illuminated with the LUYOR-3415RG Dual Fluorescent Protein Flashlight to identify the transformants. T-DNA was segregated out at T2 generation by genotyping the Cas9-encoding gene and the gene editing events were identified by PCR amplifying the targeted region followed by Sanger sequencing (Supplementary Table 1). For a mutant complementation test of *9b-1*, the fluorescence-tagged *AtALKBH9B* transgenic plants were obtained by constructing a vector *pro9B:: 9B:eGFP:terHSP18.2*. Each fragment was PCR amplified using the primers listed in Supplementary Table 1 and was cloned into the pCAMBIA1300 using the T4 DNA ligase (NEB). The construct was transformed into the *atalkbh9b-1* mutant and the transgenic lines were identified for homozygosity at T3 generation.

*Arabidopsis* seeds were surface sterilized in 75% ethanol for 15 min, washed with 100% ethanol for 1 min, and planted on half-strength MS media (including 1% sucrose). Prior to germination, seeds were stratified for 2 days at 4 °C under the dark condition and moved to a growth chamber set at 22 °C and 12-h light/12-h dark cycle. For the heat stress treatment, plants were grown for 6 days at 22 °C and then treated with heat stress of 37 °C for 24 h. To detect the retrotransposition of *Onsen*, *Arabidopsis* seedlings were grown in the media containing zebularine (Sigma) and α-amanitin (MCE), and then heat-stressed as described above. The chemical reagents were prepared by filter-sterilization (zebularine, 5 mg/ml in DMSO; α-amanitin 1 mg/ml in water) and used at the concentrations indicated in the previous study (Thieme et al., 2017). The chemical- and/or heat-stressed plants were transferred to soil and grown to maturity under the 16-h light /8-h night cycle at 22 °C.

### RT-qPCR

Plant samples were flash frozen and ground in liquid nitrogen. Total RNA was isolated using the TRIzol Universal Reagent (Tiangen). Briefly, 100 mg of the ground tissue powders were resuspended in 1 mL of TRIzol reagent, incubated at room temperature for 5 min, and then centrifuged at top speed for 10 min at 4 °C. The supernatant was mixed vigorously with chloroform and centrifuged at top speed for 10 min at 4 °C. The upper phase was mixed with the same volume of isopropanol and incubated at −80 °C for 10 min. The RNA was precipitated by centrifugation and the pellet was washed with 1 mL of 75% ethanol.

The first-strand cDNA synthesis was performed using 500 ng of RNA by the ReverTra Ace qPCR RT Master Mix with gDNA Remover (Toyobo). The resulting cDNA was diluted four-fold with DEPC-treated water and 1.5 μL was used for a 20-μL qPCR reaction mixture. The qPCR was carried out using ChamQ Universal SYBR qPCR Master Mix (Vazyme) in the CFX96 Connect Real-time PCR Detection system (BioRad). *Actin2* was used as the internal control and the sequences of the primers used for RT-qPCR is provided in Supplementary Table 1.

### RIP-qPCR

Direct binding of a protein and RNA was assessed by RIP-qPCR experiments. 7-d-old seedlings heat-stressed at 37 °C for 1 day were flash frozen and ground in liquid nitrogen. Over 2 g of frozen powder was homogenized in 6 mL of extraction buffer (100 mM Tris-HCl (pH 7.5), 150 mM NaCl, 0.5% IGEPAL (Sigma), and 1% plant protease inhibitor cocktail (MedChem Express)). The crude extract was incubated at 4 °C for 30 min with shaking and then centrifuged for 30 min at 18,000 *g* at 4 °C. 87.5 µL of 40 U/µL RNase inhibitor (ABclonal) and 25 µL of GFP-trap magnetic beads (Chromotek) was added to 3.5 mL of the supernatant and incubated overnight at 4 °C. 350 µL of the supernatant was kept as an input sample and stored at −80 °C freezer until use. After washing four times with 1 mL extraction buffer, the beads were resuspended in 150 µL of the proteinase K buffer (15 µL 10% SDS, 18 µL 10 mg/mL proteinase K and 117 µL extraction buffer) and incubated for 30 min at 55 °C. RNA was then extracted by adding 400 µL phenol:chloroform:isopropanol. The mixture was vortexed rigorously for 15 sec and centrifuged at 14,000 *g* for 10 min at room temperature. 350 μL of the aqueous phase was mixed with 400 µL of chloroform, vortexed, and centrifuged at 14,000 *g* for 10 min at room temperature. 300 μL of the aqueous phase was carefully moved to a new tube and added with 30 μL of 3 M sodium acetate (pH 5.2) and 750 μL of 100% ethanol. The mixture was incubated at −80 °C overnight and centrifuged at 14,000 *g* for 30 min at 4 °C. The pellet was washed with 80% ethanol, air-dried, and resuspended in 15 µL of RNase-free water. The input fraction was subjected to the same procedure to extract RNA. The extracted RNAs were reverse-transcribed and analyzed in qPCR as described above in RT-qPCR (the oligonucleotide sequences are provided in Supplementary Table 1). The *Onsen* LTR::*Onsen* Gag:eGFP was constructed by modifying the pGPTVII binary vector using the primers listed in Supplementary Table 1 (Gaubert et al., 2017). The construct was transformed into Col-0 and introduced to the *atalkbh9b-1* mutant by genetic cross.

The m^6^A enrichment experiment was performed as described previously with minor modifications (Dominissini et al., 2013) and the Magna RIP™ RNA-Binding Protein Immunoprecipitation Kit (Merck) was used following the manufacturer’s instruction. Briefly, 300 µg of total RNA was randomly fragmented into 250-nucleotide fragments by RNA fragmentation reagents (for 1 mL 10X reagents: 800 µL 1 M Tris-HCl (pH 7.0), 100 µL 1 M ZnCl_2_, 100 µL RNase-free H_2_O). Fragmented RNA was precipitated using 2.5 volume of EtOH, 1/10 volume of 3 M NaOH, 100 µg/mL glycogen at −80 °C overnight. After centrifugation at 14,000 *g* for 10 min, the pellet was resuspended in 55 µL RNase-free H_2_O. 5 µL of RNA was kept as the input sample and the remaining RNA was incubated with 5 µg m^6^A-specific antibody (cat. 202003, Synaptic Systems) overnight at 4 °C. The m^6^A-containing fragments were pulled down with magnetic beads. The beads were then washed five times using 500 µL of cold RIP Wash Buffer, re-suspend in 150 µL of proteinase K buffer (117 µL of RIP Wash Buffer, 15 µL of 10% SDS, 18 µL of 10 mg/mL proteinase K), and incubated at 55 °C for 30 min with shaking. After incubation, the beads were separated on magnetic rack and the supernatant was mixed with 250 µL of RIP Wash Buffer. 400 µL of phenol:chloroform:isoamyl alcohol was added and the mixture was centrifuged at 14,000 *g* for 10 min at room temperature. 350 μL of the aqueous phase was then mixed with 400 µL of chloroform, and the mixture was centrifuged at 14,000 *g* for 10 min at room temperature. 300 μL of the aqueous phase was mixed with 50 µL of Salt Solution I, 15 µL of Salt Solution II, 5 µL of Precipitate Enhancer and then 850 µL of absolute ethanol. The mixture was stored at −80 °C overnight to precipitate the RNA. Then, the mixture was centrifuged at 14,000 *g* for 30 min at 4 °C and the pellet was washed with 80% ethanol. After centrifugation at 14,000 *g* for 15 min at 4 °C, the pellet was resuspended in 20 µL of RNase-free H_2_O. The extracted RNAs were reverse-transcribed and analyzed in qPCR as described above in RT-qPCR (the oligonucleotide sequences are provided in Supplementary Table 1).

### ALE-qPCR

ALE-qPCR was performed as previously described (Cho et al., 2019; Wang et al., 2021). Genomic DNA was extracted using a DNeasy Plant Mini Kit (Qiagen) following the manufacturer’s instruction. 200 ng of genomic DNA and 1 pg of PCR-amplified *Evade* DNA was used for ligation with 0.5 µl of 40 µM adapter DNA overnight at 16 °C (the sequences are provided in Supplementary Table 1). The adapter-ligated DNA was purified by AMPure XP beads (Beckman Coulter) at a 1:0.5 ratio. In vitro transcription reactions were performed using a Standard RNA Synthesis Kit (NEB). 1 µg of purified RNA was subjected to reverse transcription using a Transcriptor First Strand cDNA Synthesis Kit (Roche) and 1 µl of RNase A/T1 (Thermo Fisher Scientific) was added to digest non-templated RNA for 30 min at 37 °C. Subsequently, qPCR was performed as described above (the oligonucleotide sequences are provided in Supplementary Table 1).

### Droplet Digital PCR

The ddPCR experiments were carried out as previously described with minor modifications (Fan and Cho, 2021). Genomic DNA was extracted using a N96 DNAsecure Plant Kit (Tiangen) following the manufacturer’s instruction. 100 ng of genomic DNA was digested using AluI for 4 h at 37 °C. The digested DNA was diluted to 0.15 ng/µL using the Qubit4 DNA quantification system (Thermo Fisher Scientific) and the Probe ddPCR SuperMix mixture was prepared (Targeting One; 15 µL 2x SuperMix, 2.4 µL (10 µM) for each primer, 0.75 µL FAM-Probe (10 µM), 0.75 µL HEX-Probe (10 µM), 3.9 µL diluted DNA totaling 30 µL). Droplets were generated using the Drop maker (Targeting One) and PCR was performed as following: 95 °C for 10 min; then 55 cycles of 94 °C for 30 sec and 56.8 °C for 30 sec; 98°C for 10 min. PCR products were read by Chip reader system (Targeting One). *CBF2* was used as the internal single-copy control. The oligonucleotide sequences are provided in Supplementary Table 1.

### Stress granule enrichment

The enrichment of cytoplasmic RNA granules was performed following the previously described method (Kim et al., 2021; Lei et al., 2021). Briefly, 2 g of seedlings was ground in liquid nitrogen and resuspended in 5 mL of lysis buffer (50 mM Tris–HCl (pH 7.4), 100 mM KOAc, 2 mM MgOAc, 0.5 mM dithiothreitol, 0.5% NP40, complete EDTA-free protease inhibitor cocktail (Roche), and 40 U/mL RNasin Plus RNase inhibitor (Promega)). The mixture was filtered through four layers of Miracloth (Sigma-Aldrich) and centrifuged at 4,000 *g* for 10 min at 4 °C. The supernatant was removed, and the pellet was resuspended in 2 mL of lysis buffer. The samples were again centrifuged at 18,000 *g* for 10 min at 4 °C. The pellet was resuspended in 2 mL of lysis buffer, vortexed and centrifuged at 18,000g at 4 °C for 10 min. The supernatant was discarded, and the pellet was resuspended in 1 mL of lysis buffer. After a brief centrifugation at 850 *g* for 10 min at 4 °C, the RNA granule fraction in the supernatant was collected for RNA extraction.

### RNA-seq analyses

The mRNAs were purified from 3 µg of total RNA using the poly-T oligo-attached magnetic beads. Library preparation was performed using the NEBNext Ultra RNA Library Prep Kit (NEB) following the manufacturer’s instructions. Sequencing was performed on an Illumina NovaSeq 6000 platform, and 150-bp paired-end (PE150) reads were generated.

For the data analysis, the raw sequences were processed using Trimmomatic (version 0.39) (Bolger et al., 2014) to remove the adapter and low-quality sequences. Trimmed reads were then aligned to the *Arabidopsis* reference genome (TAIR10) with default settings using Hisat2 (version 2.2.1) (Kim et al., 2015). The FPKM values of genes and TEs were calculated by StringTie (version 2.1.7) (Pertea et al., 2015). Visualization of the sequencing data was performed using the Integrative Genomics Viewer (IGV) (Robinson et al., 2011). For the m^6^A peak calling, MACS2 (version 2.2.7.1) (Zhang et al., 2008) was run with the following parameters; --nomodel,--extsize 50, -p 5e-2, and -g 65084214 (the -g option accounts for the size of the *Arabidopsis* transcriptome).

### Confocal microscopy

To determine the subcellular localization of SGS3 and AtALKBH9B proteins, the pGPTVII binary vector was modified to generate *proUBQ10::SGS3:TdTomato* and *proUBQ10::9B:eGFP* constructs. The CDS of *AtALKBH9B* and *SGS3* was amplified from Col-0 cDNA using KOD-Plus-Neo (Toyobo) (primers are listed in Supplementary Table S1). The vectors were transformed into Col-0 plants, which were selected on 1/2 MS plates containing 10 μg/mL Glufosinate ammonium (Coolaber) and further confirmed by PCR using the primers targeting *GFP* and *TdTomato* (listed in Supplementary Table 1). The *9B-GFP SGS3-TdTomato* double transgenic line was generated from genetic crossing and subsequent PCR-based genotyping in F2 populations. The transgenic plants were heat-stressed at 37 °C for 12 h and the fluorescence signals were detected by Zeiss LSM880 confocal microscopy. For the tobacco transient expression experiments, the constructs were expressed with P19 in tobacco leaves. Tobacco plants were heat-stressed at 37 °C for 12 h at 48 h after agro-infiltration.

### IP-MS

The 7-d-old seedlings of *proUBQ10::AtALKBH9B:GFP* and *pro35S::GFP* were treated with 1-d heat stress under 37 °C, and immediately flash frozen. 1 g of ground powder was homogenized in 3 mL of IP buffer (20 mM HEPES (pH 7.4), 2 mM EDTA, 25 mM NaF, 1 mM Na_3_VO_4_, 10% Glycerol, 100 mM NaCl, 0.5% Triton X -100 and 1% plant protease inhibitor cocktail (MedChem Express)) and the mixture was rotated at 4 °C for 1 h. The crude extract was centrifuged for 20 min at 18,000 *g* at 4 °C. The immunoprecipitation was performed using 3 mL of plant extract mixed with 25 µL of GFP-trap magnetic beads (Chromotek) at 4 °C overnight. The beads were washed four times with 1 mL IP buffer and centrifuged for 1 min at 200 *g* at 4 °C.

For protein digestion, 100 μg of protein was reduced with 2 µl 0.5 M Tris(2-carboxyethyl)phosphine (TCEP) at 37 °C for 60 min and alkylated with 4 µl 1 M iodoacetamide (IAM) at room temperature for 40 min in darkness. Five-fold volumes of cold acetone were added to precipitate protein at −20 °C overnight. After centrifugation at 12,000 *g* at 4 °C for 20 min, the pellet was washed twice using 1 mL pre-chilled 90% acetone aqueous solution. Then, the pellet was re-suspended with 100 µl 10 mM Triethylammonium bicarbonate (TEAB) buffer. Trypsin (Promega) was added at 1:50 trypsin-to-protein mass ratio and incubated at 37 °C overnight. The peptide mixture was desalted by C18 ZipTip, and lyophilized by SpeedVac.

For nano-HPLC-MS/MS analysis, the peptides were analyzed by online nano flow liquid chromatography tandem mass spectrometry performed on an EASY-nanoLC 1200 system (Thermo Fisher Scientific) connected to a Q Exactive™ Plus mass spectrometer (Thermo Fisher Scientific). Acclaim PepMap C18 (75 μm x 25 cm) was equilibrated with solvent A (A: 0.1% formic acid in water) and solvent B (B: 0.1% formic acid in ACN). 3 μL peptide was loaded and separated with 60 min-gradient at flow rate of 300 nL/min. The column temperature was 40 °C. The electrospray voltage of 2 kV versus the inlet of the mass spectrometer was used. The peptides were eluted using the following gradient: 0-3 min, 2-6% B; 3-42 min, 6−20% B; 42-47 min, 20-35% B; 47-48 min, 35-100% B; 48-60 min, maintained 100% B.

The mass spectrometer was run under data dependent acquisition (DDA) mode, and automatically switched between MS and MS/MS mode. The survey of full scan MS spectra (m/z 200-1800) was acquired in the Orbitrap with resolution of 70,000. The automatic gain control (AGC) target at 3e6 and the maximum injection time was 50 ms. Then, the top 20 most intense precursor ions were selected into collision cell for fragmentation by higher-energy collision dissociation (HCD) with the collection energy of 28. The MS/MS resolution was set at 17500, the automatic gain control (AGC) target at 1e5, the maximum injection time was 45 ms, isolation window was 2 m/z, and dynamic exclusion was 30 seconds.

Tandem mass spectra were processed by PEAKS Studio (v.10.6, Bioinformatics Solutions Inc.). PEAKS DB was set up to search the uniprot_Arabidopsis_thaliana (v.201907, entries 27477) database assuming trypsin as the digestion enzyme. PEAKS DB was searched with a fragment ion mass tolerance of 0.02 Da and a parent ion tolerance of 7 ppm. Carbamidomethylation (C) was specified as the fixed modification. Oxidation (M), Deamidation (NQ), and Acetylation (K) were specified as the variable modifications. The peptides with −10logP≥20 and the proteins with −10logP≥20 containing at least 1 unique peptide were filtered.

### Split luciferase complementation assay

The *AtALKBH9B* and *ECT2* were amplified by PCR and cloned into the modified pCAMBIA_nLUC and pCAMBIA_cLUC vectors containing the 35S promoter (primers are listed in Supplementary Table 1). The *pro35S::9B:nLUC* and *pro35S::cLUC:ECT2* constructs were transformed into the *Agrobacterium tumefaciens* strain GV3101 and then infiltrated into *Nicotiana benthamiana* leaves along with P19. The detached leaves were sprayed with 1 mM luciferin (GLPBio) at 2 days after infiltration. The luminescence signal was visualized with a Tanon-5200 (Tanon), and the images were acquired and processed using AllCap software (Tanon).

### Fluorescence polarization (FP)

The Gag region of *Onsen* was PCR-amplified using the primers listed in Supplementary Table 1, cloned into pET28a generating 6xHis:*Onsen* Gag and transformed into *E. coli* strain Rosetta. Starter culture was grown overnight in 4 mL LB media containing 50 µg/mL Kanamycin and 25 µg/mL Chloramphenicol at 37 °C with shaking at 200 RPM. 3 mL of starter culture was transferred to 300 mL of LB media. Cells were grown at 37 °C with shaking at 200 RPM until the OD600 reach between 0.6 and 0.8. The growth temperature was then lowered to 12 °C and IPTG was added to a final concentration of 0.5 mM. Cells were incubated for two days at 12 °C with shaking at 180 RPM and harvested by centrifugation. The pellet was resuspended in 30 mL of lysis buffer (20 mM Tris-HCl (pH 7.6), 200 mM NaCl, 10% Glycerol, 0.1% Tween20). 60 µL 1 U/µL DNase I, 60 µL 1 M MgSO_4_ and 150 µL 200 mM PMSF were added, and the cells were lysed using an SCIENTZ-IID cell homogenizer (SCIENTZ). Lysates were cleared with cell debris by centrifugation at 18,000 *g* for 1 h at 4 °C. Cleared lysates were loaded onto a Econo-Pac® Chromatography Columns column (Bio-rad) and washed with 2 column volumes of buffer (20 mM Tris-HCl (pH 7.6), 200 mM NaCl, 10% Glycerol, 0.1% Tween20, 25 mM imidazole). Bound protein was eluted in 2 mL of buffer (20 mM Tris-HCl (pH 7.6), 200 mM NaCl, 10% Glycerol, 0.1% Tween20, 500 mM imidazole). Protein was concentrated using a spin concentrator (Amicon, 10K MWCO) and injected onto Superdex 200 column (GE Healthcare) equilibrated in 25 mM HEPES (pH 7.5) and 100 mM NaCl. Fractions were checked for purity by SDS-PAGE followed by Coomassie blue staining. Fluorescence polarization assay was carried out following the previously described method (Harrison et al., 2016). Binding assays were performed in 25 mM HEPES (pH 7.5) and 100 mM NaCl including 10 nM FAM-labeled RNA oligonucleotide (GGCCAACUACGU and GGCCAm^6^ACUACGU) in black and flat-bottom 96-well plates (BBI). Proteins were serially diluted 2-fold and the final assay volume was 25 µL per well. The signal was detected at room temperature on a BioTek Synergy Neo plate reader (BioTek). Polarization (P) was converted to anisotropy (A) using the formula A = 2P/(3-P). Data were plotted as fraction bound by setting the highest anisotropy measured to 1. Data were plotted using GraphPad Prism (version 6.0), and dissociation constants (*Kd*) were obtained by fitting the curve to a non-linear regression model.

### Reverse transcription efficiency

RT efficiency assay was performed using the MEGAscript® RNAi Kit (Thermo Fischer) according to the manufacturer’s instructions. In brief, the template DNA was amplified using the primers containing T7 promoter (listed in the Supplementary Table 1). The in vitro transcription was carried out as following; 3 µL 10X T7 Reaction Buffer, 0.5 µL ATP Solution or 0.5 µL m^6^ATP (TriLink), 1 µL each of C/G/UTP, 1 µL T7 RNA polymerase, 2.5 µL DNA (500 ng) at 37 °C overnight. The template DNA was removed by adding the DNase I and incubating the mixture at 37 °C for 2 h. RNA was purified by the ethanol precipitation method and resuspended in 20 µL RNase-free H_2_O. RNA was reverse transcribed using the ReverTra Ace qPCR RT Master Mix with gDNA Remover (Toyobo) for a limited time period as indicated in the figure legends. Subsequently, q PCR was performed as above described. The oligonucleotide sequences are provided in Supplementary Table 1.

### Data availability

The NGS data generated in this study are deposited to SRA repository under PRJNA793364 and summarized in Supplementary Table 2. The analyses were performed using the standard codes instructed by the tools described in the Methods and the custom codes used in this study are deposited to GitHub (https://github.com/JungnamChoLab).

## Acknowledgements

We thank the Core Facility Center of CAS Center for Excellence in Molecular Plant Sciences for technical support with confocal microscopy. This work was supported by the Strategic Priority Research Program of the Chinese Academy of Sciences (XDB27030209), the National Natural Science Foundation of China (31970518, 32150610473 and 32111540256) and General Program of Natural Science Foundation of Shanghai (22ZR1469100).

## Author contributions

WF and JuC conceived the idea and designed the experiments. WF, ZL and JiC conducted the experiments. WF and LW analyzed the data. WF and JuC drafted the manuscript. JuC revised the manuscript.

