## Supplementary information for "Suppression of transposon mobilization by m^6^A-mediated RNA sequestration in stress granules"

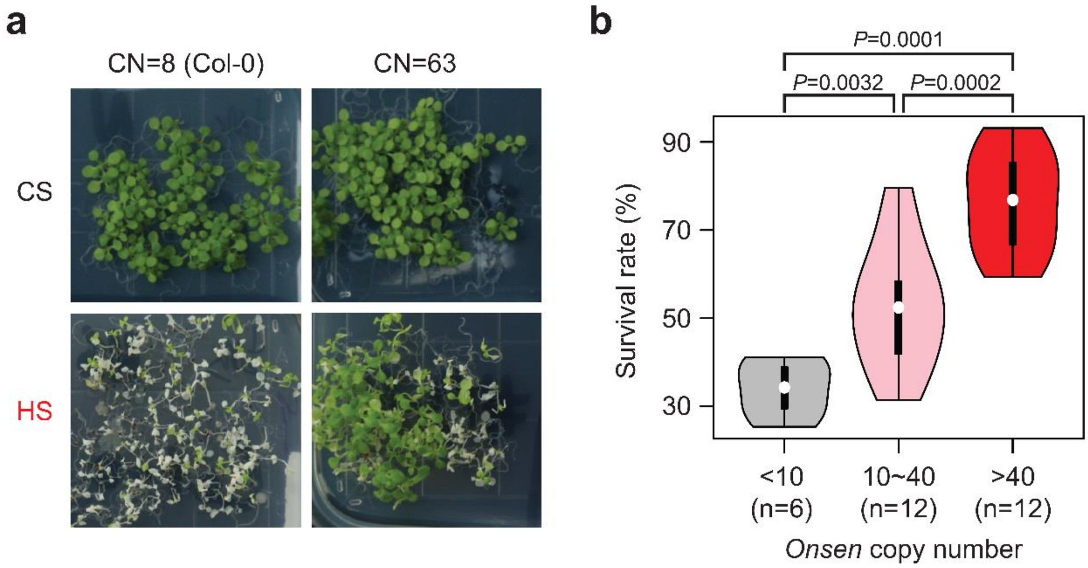

**Supplementary Fig. 1 | Increased heat tolerance of *Onsen*-proliferated *Arabidopsis* plants.**

**a**. Representative images of *Arabidopsis* plants containing different copy number of *Onsen*. CN, copy number; CS, control sample; HS, heat-stressed sample. **b**. Survival rate after 24 h of heat treatment at 37 °C. Plant lines were grouped by their *Onsen* copy numbers. Number of plant lines used are shown in parentheses. Open circle indicates median value and black rectangle represents interquartile range. *P* values were obtained by the one-sided Wilcoxon rank sum test.

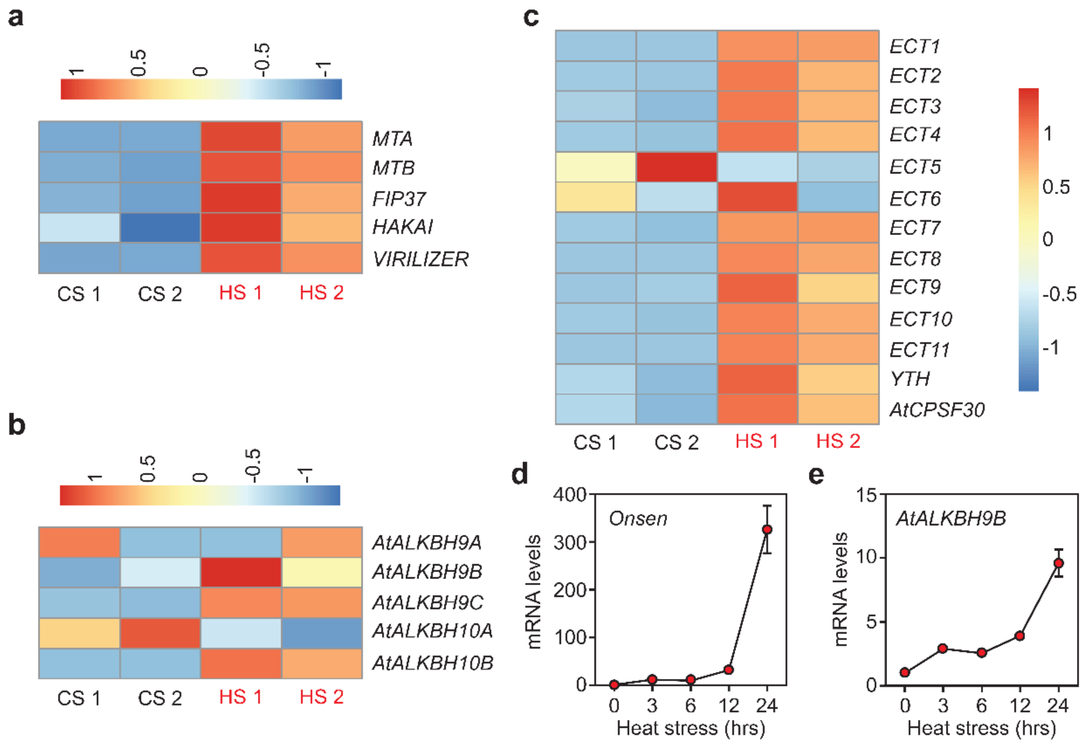

**Supplementary Fig. 2 | Heat responsiveness of m$^6$A-related factors.**

**a**-**c**. Heatmap displaying the expression profiles of m$^6$A methyltransferases (**a**), demethylases (**b**), and readers (**c**). **d** and **e**. RT-qPCR validation for the expression pattern of *Onsen* (**d**) and *AtALKBH9B* (**e**) in various duration of the heat stress treatment. Values are mean ± s.d. from three biological replications.

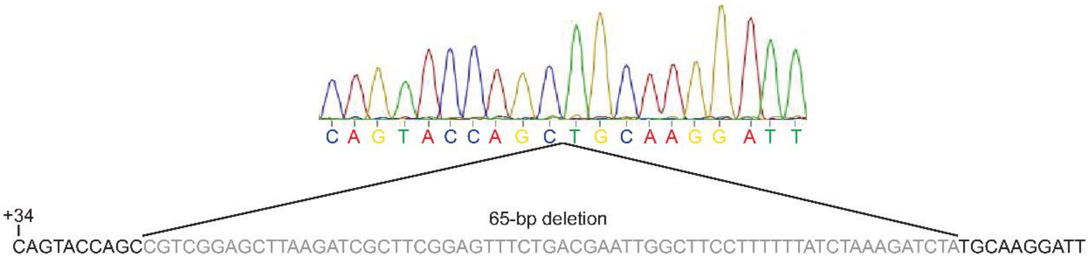

**Supplementary Fig. 3 | *De novo* mutation of *AtALKBH9B* by CRISPR-Cas9.**

Chromatogram of Sanger sequencing results for the *9b-2* mutant. *Arabidopsis* plants transformed with CRISPR-Cas9 construct were screened at T2 generation.

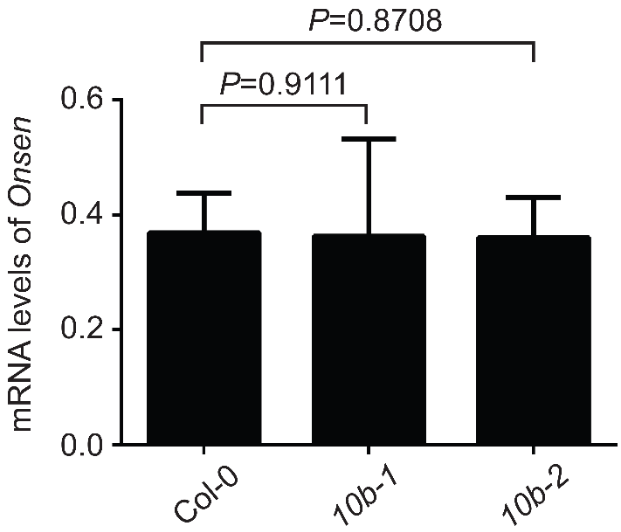

**Supplementary Fig. 4 | *Onsen* mRNA levels in the mutants of *AtALKBH10B*.**

The expression levels of *Onsen* in the *10b-1* and *10b-2* mutants determined by RT-qPCR. Plants were heat-stressed at 37 °C for 24 h and harvested for RNA extraction immediately after the heat treatment. Values are mean ± s.d. from three biological replications.

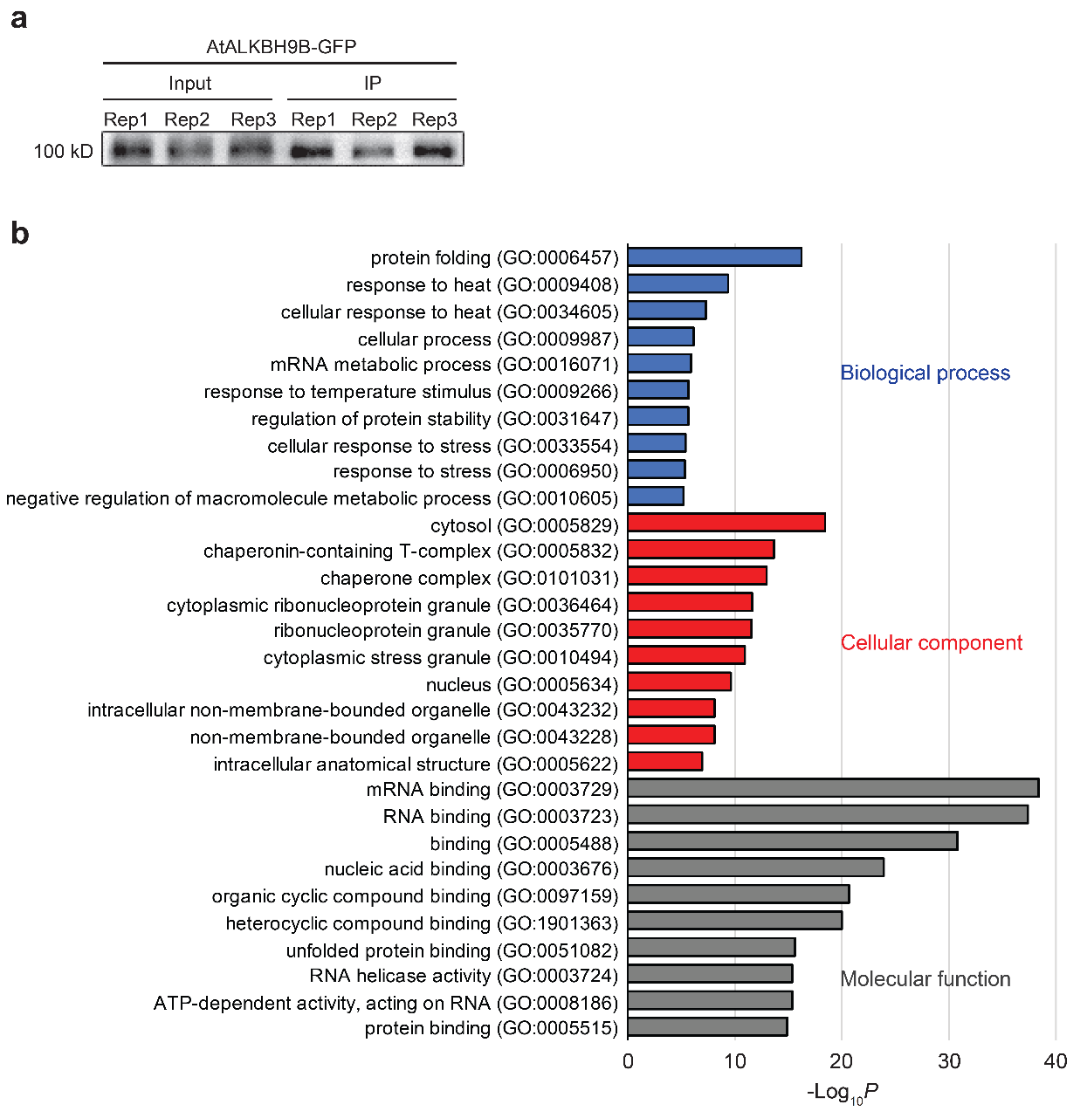

**Supplementary Fig. 5 | AtALKBH9B interactome.**

**a**. Western blot image showing 9B-GFP in the input and immunoprecipitated samples.

**b**. Gene enrichment analysis of the proteins identified by the IP/MS experiment using the 9B-GFP transgenic plants.

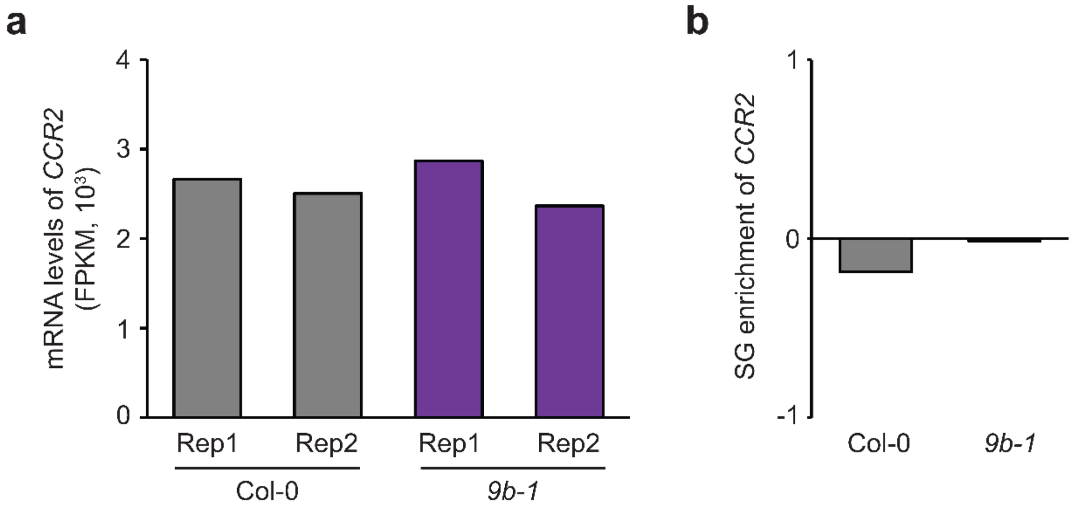

**Supplementary Fig. 6 | SG enrichment and m$^6$A level of *CCR2*.**

**a**. The mRNA levels of *CCR2* in the wt and *9b-1* mutants assessed by RNA-seq. **b**. The SG enrichment score of *CCR2* in the wt and *9b-1* mutants. SG enrichment score was determined by log2-transformed fold change of SG fraction levels to total RNA levels.

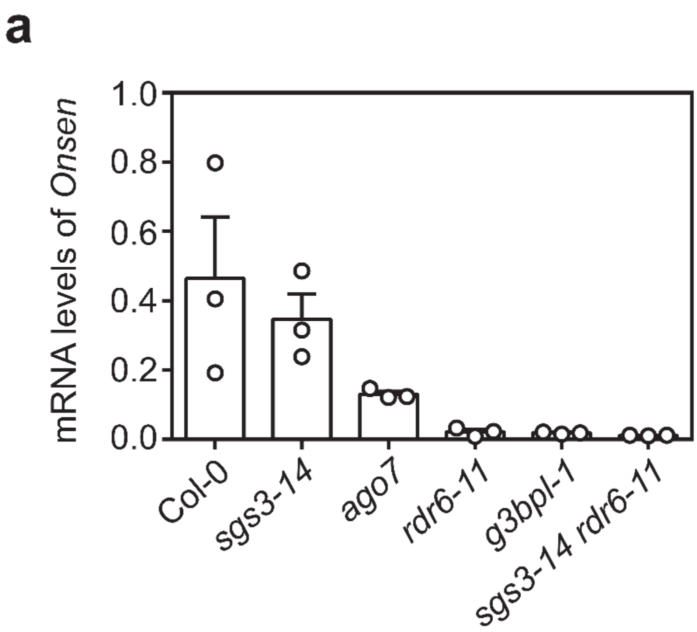

**Supplementary Fig. 7 | *Onsen* RNA levels in the mutants of SG components.**

**a**. The levels of *Onsen* RNA determined by RT-qPCR. Plants were heat-stressed for 24 h at 37 °C and harvested immediately after the heat treatment. Values are mean ± s.d. from three biological replications.

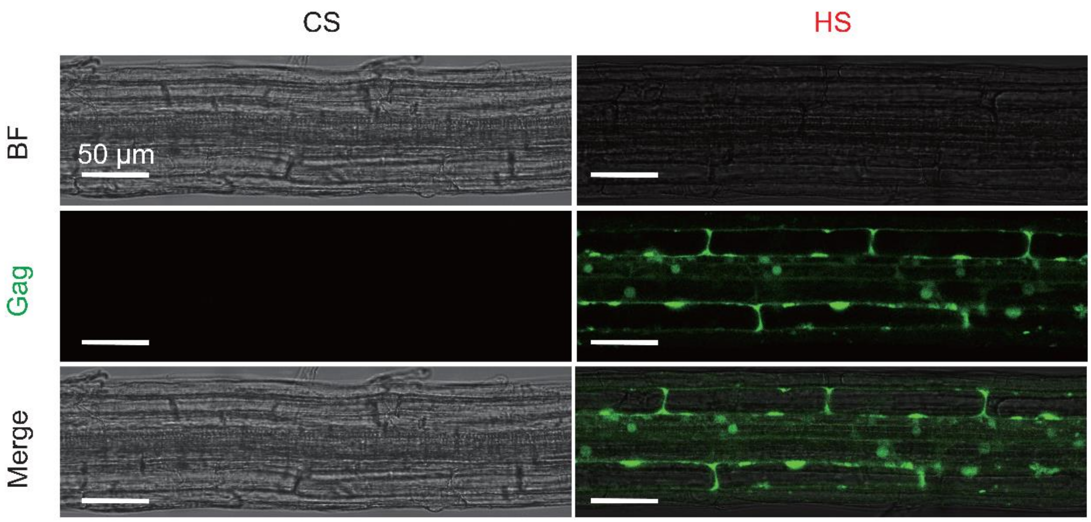

**Supplementary Fig. 8 | Confocal microscopy images of Gag-GFP transgenic plants.** Representative confocal microscopy images of *Arabidopsis* transgenic plants containing the *OnsenLTR::Gag:GFP* construct. Plants were tested for green fluorescence in control (CS) and heat stress (HS) conditions with pre-heating for 12 h at 37 °C before microscopy imaging. Bar=50 µm.

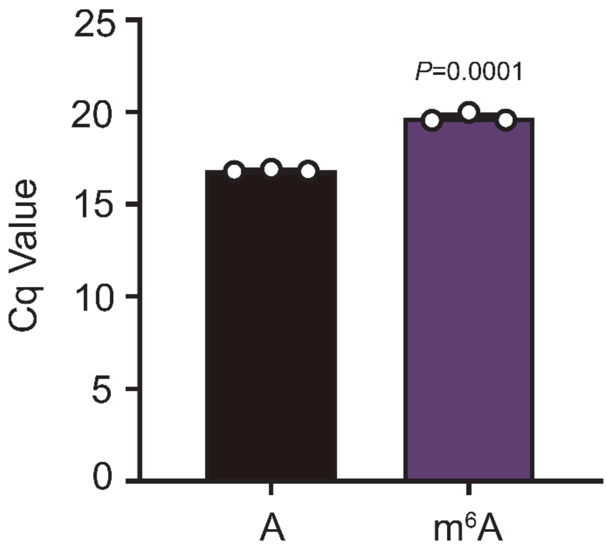

**Supplementary Fig. 9 | Reverse transcription efficiency of m6A-modified RNA.**

RT-qPCR experiments using the in vitro transcribed *Onsen* RNAs that include either A or m6A. Values are mean ± s.d. from three biological replications. *P* values were obtained by the one-sided Student’s t test.

**Supplementary Table 1 | Oligonucleotide sequences used in this study.**

| Primer name | Sequence（5' to 3'） |
|---|---|
| **Genotype** | |
| *9b-1*_genotyping-F | CGAGTTCGATGAAGACTCCAG |
| *9b-1*_genotyping-R | ATCCTGTTGAATAGAACCGGG |
| *9b-2*_genotyping-F | AGAATTTTAAACGGCCCAAGAG |
| *9b-2*_genotyping-R | AGCTCTTTCTGATCCCATATTTTCC |
| *10b-1*_genotyping-F | TCCCTCTCATCACCAACAAAG |
| *10b-1*_genotyping-R | ATGCCATAGCCATGAAGATTG |
| *10b-2*_genotyping-F | AGTAGAAAACACATGCCTCGG |
| *10b-2*_genotyping-R | TTAACATCGAGCCAATTCCAC |
| *ago7-1*_genotyping-F | GTATTCTGGAGGCAGAGGAGC |
| *ago7-1*_genotyping-R | CTCCTCCTTTCTTTTGCACC |
| *sgs3-14*_genotyping-F | AAATTTGGAGTCCAGAATCGG |
| *sgs3-14*_genotyping-R | CAAAGCATCGGAATCATTCTC |
| *g3bpl-1*_genotyping-F | AAATGACAAGACCGGATCATG |
| *g3bpl-1*_genotyping-R | TATCAAGTGTTGCAGCAGCAG |
| *Cas9*_genotyping-F | TCCACACCTGAAGCGTTGATAG |
| *Cas9*_genotyping-R | ATGGATAAGAAGTACTCTATCGGACT |
| *eGFP*_genotyping-F | GTGAACCGCATCGAGCTGAA |
| *eGFP*_genotyping-R | ACGTTGTGGCTGTTGTAGTTG |
| *TdTomato*_genotyping-F | TAATGCAGAAGAAGACCATGGGC |
| *TdTomato*_genotyping-R | GGAAGGACAGCTTCTTGTAATC |
| SALK_LB1.3 | ATTTTGCCGATTTCGGAAC |
| **Cloning** | |
| *9B* promoter_EcoRI-F | CGCGAATTCTGTGGTGTGGTGTGGTGTG |
| *9B* promoter_SacI-R | CGGGAGCTCGAGATACGCTCGATACAATCCAAA |
| *9B* gDNA_SacI-F | CGGGAGCTCATGGAAAACGATCCATTTCTCCG |
| *9B* gDNA_KpnI-R | TGCGGTACCACCGTAGTTTCTTCTACTAGGACG |
| *eGFP*_XbaI-F | TGCTCTAGAATGGTGAGCAAGGGCGAG |
| *eGFP*_PstI-R | CGCCTGCAGCTTGTACAGCTCGTCCATGC |
| *HSP18.2*_PstI-F | CGCCTGCAGTGAATATGAAGATGAAGATGAAATATTT |
| *HSP18.2*_HindIII-R | CGCAAGCTTCTTATCTTTAATCATATTCCATA |
| *Onsen* LTR_HindIII-F | CGCAAGCTTTGTTGAAAGTTAAACTTGATTTTG |
| *Onsen* LTR_BamHI-R | CGCGGATCCTGTTAGAGTAAAATTCTTTTAGAG |
| *Onsen* Gag_BamHI-F | CGCGGATCCATGAGAGACTCAAGAAAGAGAGAC |
| *Onsen* Gag_XhoI-R | CGCCTCGAGATCTTCTTTCTTCTTCTTCTTTTC |
| *9B* CDS_SpeI-F | CGGACTAGTATGGAAAACGATCCATTTCTCCG |
| *9B* CDS_XhoI-R | CCGCTCGAGACCGTAGTTTCTTCTACTAGGACG |
| *9b*_sgRNA1-top | ATTGTCTCCGGCAGTACCAGCCGT |
| *9b*_sgRNA1-bottom | AAACACGGCTGGTACTGCCGGAGA |
| *9b*_sgRNA2-top | ATTGCTTCGGAGTTTCTGACGAAT |

| | |
|---|---|
| *9b*_sgRNA2-bottom | AAAGATTCGTCAGAAACTCCGAAG |
| *9b*_sgRNA3-top | ATTGTTTATCTAAAGATCTATGCA |
| *9b*_sgRNA3-bottom | AAAGTGCATAGATCTTTAGATAAA |
| pET28a_*Onsen* Gag-F | ACAGCAAATGGGTCGCATGAGAGACTCAAGAAAG |
| pET28a_*Onsen* Gag-R | CGACGGAGCTCGAATTCGGATCCATCTTCTTTCTTCTTC |
| JW771_9B-F | TTTGGAGAGAACACGGGGGACATGGAAAACGATCCATT |
| JW771_9B-R | TAGTCCATTTGTTGGATCCCGACCGTAGTTTCTTCTACTA |
| JW772_ECT2-F | CGAGAAGCTCGAGTATCTTTTAAAACAAAAGAGGA |
| JW772_ECT2-R | GATACGAACGAAAGCTCGGCAAGATAGATCAAAA |
| **In vitro Transcription** | |
| *Onsen* fragment PCR-F | TAATACGACTCACTATAGGGCCACCATGCTCCTAGCAACAAAAAATTTGAG |
| *Onsen* fragment PCR-R | ATCGAATGAGAATGTTTCCTTTACCT |
| **ddPCR** | |
| *Onsen* ddPCR-F | GAAAAGAAGAAGAAGAAAGAAGATAT |
| *Onsen* ddPCR-R | CCATTTCCATATCCACCACG |
| *CBF2* ddPCR-F | CTTCGGCCATGTTATCCAAC |
| *CBF2* ddPCR-R | TTTATACGCCGGAACAGAGC |
| *Onsen* ddPCR-Probe | TAGACATCCCCAACATCGCCTCTTCAT (5'-HEX, 3'-BHQ1) |
| *CBF2* ddPCR-Probe | CCAAAGTTACCAAAGAAGAGGTGGTGGT (5'-FAM, 3'-BHQ1) |
| **ALE-qPCR** | |
| Adaptor_ALE-top | AGAGAGTAATACGACTCACTATAGGGACACGACGCTCTTCCGATCT |
| Adaptor_ALE-bottom | AGATCGGAAGAGCGTCGTGTCCCTATAGTGAGTCGTATTACTCTCT (5'-phos) |
| ALE RT primer | AGACGTGTGCTCTTCCGATCTGCTCTGATACCA |
| *Evade* full length-F | TATTGATCAAGACTCAAATAAGAAAG |
| *Evade* full length-R | AAGAGTGAGATAGATCCACAAG |
| *Onsen* ALE_qPCR-F | CCGATCTTGTTGAAAGTTAAACT |
| *Onsen* ALE_qPCR-R | TCTAGAACTTGGATTTGGCC |
| *Evade* ALE_qPCR-F | TCCGATCTTATTGATCAAGAC |
| *Evade* ALE_qPCR-R | AGACTTCTCATATGTTCGGC |
| **Quantitative PCR** | |
| *Actin2*_qPCR-F | GGTAACATTGTGCTCAGTGGTGG |
| *Actin2*_qPCR-R | CAACGACCTTAATCTTCATGCTGC |
| *CCR2*_qPCR-F | CGTCCGGTGATGTTGAGTATCG |
| *CCR2*_qPCR-R | TCTTGGAATCAATAACGTCGCCG |
| $m^6A$_region1_qPCR-F | TTAAAATATTTTAGATATTTTGTAGTT |
| $m^6A$ _region1_qPCR-R | TTAAGTGTTTTGAGAGAGTTTTTT |
| $m^6A$ _region2_qPCR-F | CAAGTGTCAAATGCTACAATTGTG |

| | |
|---|---|
| $m^6A$ _region2_qPCR-R | CTCAAATTTTTTGTTGCTAGGAG |
| $m^6A$ _region3_qPCR-F | GAGAAGGCCAACTACGTTGAA |
| $m^6A$ _region3_qPCR-R | ACCACTTATGATTCTCTTTTTGTTC |

**Supplementary Table 2 | Summary of NGS data.**

| Sample | Clean Reads | Uniquely mapped | Multiple mapped | Alignment rate (%) |
|---|---|---|---|---|
| $m^6$A-RIP-seq of wt CS IP rep1 | 11430584 | 9768775 | 475845 | 92.34 |
| $m^6$A-RIP-seq of wt CS Input rep1 | 18667778 | 15343391 | 570399 | 96.87 |
| $m^6$A-RIP-seq of wt CS IP rep2 | 25373399 | 21785294 | 1072423 | 92.33 |
| $m^6$A-RIP-seq of wt CS Input rep2 | 27382097 | 24132541 | 888125 | 96.32 |
| $m^6$A-RIP-seq of wt HS IP rep1 | 15847607 | 13397892 | 1027114 | 93.65 |
| $m^6$A-RIP-seq of wt HS Input rep1 | 18687601 | 15195099 | 1066624 | 97.13 |
| $m^6$A-RIP-seq of wt HS IP rep2 | 30905424 | 26232687 | 2025496 | 93.65 |
| $m^6$A-RIP-seq of wt HS Input rep2 | 36097518 | 33142202 | 1788580 | 98.70 |
| RNA-seq HS Col-0 rep 1 | 22556913 | 20281181 | 970841 | 97.46 |
| RNA-seq HS Col-0 rep 2 | 22310003 | 19866134 | 844322 | 96.08 |
| RNA-seq HS *9b-1* rep 1 | 22619925 | 19353102 | 1557708 | 96.77 |
| RNA-seq HS *9b-1* rep 2 | 21984648 | 19663770 | 1069017 | 97.78 |
| SG-RNA-seq HS Col-0 rep 1 | 20591865 | 17327121 | 1054539 | 92.49 |
| SG-RNA-seq HS Col-0 rep 2 | 22868648 | 20346404 | 1145541 | 97.08 |
| SG-RNA-seq HS *9b-1* rep 1 | 22575022 | 19845031 | 1323148 | 97.36 |
| SG-RNA-seq HS *9b-1* rep 2 | 22236087 | 20127837 | 934226 | 97.80 |

**Supplementary Table 3 | List of interacting proteins of 9B.**

Number of peptides and *P* values are shown.

| Gene id | *GFP* | *9B:GFP* rep1 | *9B:GFP* rep2 | *9B:GFP* rep3 | $-10\log_{10}P$ |
|---|---|---|---|---|---|
| AT1G02500 | 2 | 13 | 12 | 14 | 211.88 |
| AT1G07360 | 0 | 8 | 7 | 13 | 167.37 |
| AT1G10170 | 1 | 24 | 29 | 24 | 240.89 |
| AT1G10200 | 0 | 2 | 1 | 1 | 44.25 |
| AT1G11650 | 0 | 2 | 1 | 2 | 73.33 |
| AT1G16030 | 14 | 38 | 48 | 40 | 286.83 |
| AT1G17370 | 0 | 1 | 1 | 1 | 31.65 |
| AT1G20960 | 1 | 6 | 5 | 3 | 100.15 |
| AT1G24510 | 0 | 6 | 8 | 7 | 145.9 |
| AT1G29250 | 0 | 5 | 4 | 3 | 90.97 |
| AT1G30070 | 0 | 1 | 3 | 1 | 37.43 |
| AT1G33680 | 0 | 5 | 3 | 5 | 94.35 |
| AT1G43850 | 0 | 4 | 1 | 3 | 87.7 |
| AT1G45201 | 0 | 5 | 3 | 6 | 113.09 |
| AT1G47490 | 0 | 3 | 2 | 2 | 73.11 |
| AT1G47500 | 0 | 3 | 2 | 2 | 73.11 |
| AT1G48410 | 0 | 5 | 5 | 6 | 130.34 |
| AT1G49600 | 0 | 2 | 1 | 2 | 60.73 |
| AT1G49760 | 1 | 3 | 4 | 7 | 132.33 |
| AT1G66260 | 0 | 6 | 5 | 9 | 135.56 |
| AT1G72150 | 0 | 4 | 5 | 5 | 134.83 |
| AT1G72610 | 0 | 1 | 5 | 2 | 85.91 |
| AT1G75560 | 0 | 4 | 0 | 5 | 114.23 |
| AT1G75660 | 1 | 8 | 7 | 5 | 134.07 |
| AT1G76010 | 1 | 7 | 9 | 8 | 162.02 |
| AT1G79920 | 5 | 20 | 22 | 27 | 209.82 |
| AT1G80070 | 0 | 7 | 9 | 8 | 141.95 |
| AT1G80410 | 0 | 1 | 3 | 5 | 107.86 |
| AT2G02160 | 0 | 2 | 3 | 1 | 117.01 |
| AT2G17870 | 0 | 4 | 2 | 3 | 133.26 |
| AT2G17970 | 0 | 228 | 212 | 221 | 442 |
| AT2G18510 | 0 | 1 | 1 | 1 | 61.89 |
| AT2G20190 | 0 | 10 | 7 | 11 | 203.08 |
| AT2G21130 | 0 | 1 | 1 | 1 | 38.82 |
| AT2G23350 | 3 | 10 | 10 | 9 | 184.24 |
| AT2G26150 | 0 | 5 | 5 | 5 | 113.08 |
| AT2G26280 | 0 | 2 | 3 | 1 | 58.3 |
| AT2G27100 | 1 | 14 | 9 | 13 | 243.15 |
| AT2G29190 | 0 | 3 | 2 | 3 | 97.59 |

| | | | | | |
|---|---|---|---|---|---|
| AT2G32120 | 2 | 7 | 6 | 5 | 142.87 |
| AT2G32700 | 0 | 5 | 4 | 3 | 179.59 |
| AT2G33730 | 0 | 2 | 1 | 2 | 75.81 |
| AT2G36880 | 3 | 15 | 12 | 16 | 222.41 |
| AT2G42270 | 0 | 8 | 7 | 3 | 104.6 |
| AT2G42520 | 0 | 9 | 6 | 8 | 181.9 |
| AT2G45620 | 0 | 2 | 2 | 2 | 84.65 |
| AT2G45810 | 0 | 3 | 2 | 3 | 110.49 |
| AT3G01090 | 0 | 6 | 5 | 5 | 119.38 |
| AT3G01540 | 0 | 1 | 0 | 3 | 65.65 |
| AT3G02530 | 0 | 12 | 7 | 11 | 178.44 |
| AT3G03060 | 0 | 1 | 2 | 2 | 81.59 |
| AT3G03950 | 0 | 4 | 4 | 2 | 138.52 |
| AT3G03960 | 0 | 6 | 6 | 4 | 162.61 |
| AT3G04590 | 0 | 2 | 1 | 3 | 88.66 |
| AT3G04610 | 1 | 5 | 3 | 4 | 119.78 |
| AT3G06410 | 0 | 1 | 1 | 1 | 40.4 |
| AT3G06480 | 0 | 2 | 1 | 3 | 90.17 |
| AT3G09440 | 9 | 37 | 28 | 34 | 263.77 |
| AT3G09840 | 0 | 1 | 1 | 1 | 61.78 |
| AT3G11830 | 0 | 8 | 11 | 5 | 186.7 |
| AT3G11910 | 0 | 5 | 6 | 5 | 143.48 |
| AT3G12050 | 0 | 2 | 1 | 3 | 102.44 |
| AT3G12130 | 1 | 5 | 3 | 5 | 116.02 |
| AT3G13300 | 0 | 1 | 2 | 0 | 62.62 |
| AT3G13460 | 0 | 3 | 2 | 3 | 101.76 |
| AT3G14100 | 0 | 1 | 1 | 1 | 31.65 |
| AT3G15010 | 0 | 2 | 2 | 3 | 93.95 |
| AT3G16420 | 0 | 1 | 3 | 1 | 99.97 |
| AT3G17390 | 2 | 17 | 14 | 17 | 223.21 |
| AT3G18190 | 0 | 7 | 10 | 5 | 155.74 |
| AT3G19130 | 0 | 2 | 1 | 2 | 60.73 |
| AT3G20050 | 0 | 4 | 9 | 8 | 168.79 |
| AT3G23300 | 0 | 4 | 3 | 1 | 91.15 |
| AT3G27700 | 0 | 1 | 1 | 1 | 46.67 |
| AT3G29160 | 0 | 3 | 1 | 2 | 81.65 |
| AT3G50670 | 1 | 3 | 3 | 8 | 116.09 |
| AT3G53110 | 0 | 2 | 1 | 1 | 60.76 |
| AT3G53520 | 0 | 1 | 2 | 5 | 103.76 |
| AT3G54470 | 0 | 3 | 3 | 1 | 116.26 |
| AT3G58510 | 0 | 10 | 10 | 11 | 196.36 |
| AT3G58570 | 0 | 6 | 5 | 6 | 150.37 |
| AT3G59350 | 0 | 4 | 2 | 4 | 104.45 |
| AT3G61240 | 0 | 5 | 3 | 5 | 143.97 |

| | | | | | |
|---|---|---|---|---|---|
| AT3G62830 | 0 | 1 | 4 | 3 | 118.15 |
| AT4G00660 | 0 | 7 | 1 | 6 | 125.2 |
| AT4G01850 | 1 | 12 | 9 | 13 | 207.11 |
| AT4G03110 | 0 | 2 | 2 | 1 | 48.56 |
| AT4G08350 | 0 | 6 | 1 | 1 | 86.62 |
| AT4G09150 | 1 | 12 | 14 | 14 | 153.65 |
| AT4G14360 | 0 | 3 | 2 | 2 | 89.1 |
| AT4G16830 | 0 | 4 | 1 | 1 | 62.74 |
| AT4G23650 | 0 | 1 | 1 | 2 | 81.17 |
| AT4G27320 | 1 | 3 | 4 | 5 | 138.14 |
| AT4G31770 | 0 | 4 | 1 | 1 | 95.52 |
| AT4G34110 | 2 | 6 | 7 | 9 | 160.5 |
| AT4G34660 | 0 | 2 | 2 | 3 | 98.14 |
| AT4G38130 | 0 | 2 | 1 | 1 | 58.01 |
| AT4G38740 | 0 | 1 | 1 | 1 | 38.82 |
| AT5G02530 | 0 | 9 | 7 | 10 | 161.84 |
| AT5G03280 | 0 | 2 | 3 | 3 | 124.39 |
| AT5G03340 | 0 | 1 | 1 | 1 | 61.78 |
| AT5G06600 | 0 | 9 | 10 | 7 | 167.11 |
| AT5G08450 | 0 | 5 | 4 | 4 | 106 |
| AT5G09880 | 0 | 2 | 1 | 1 | 83.23 |
| AT5G13010 | 0 | 9 | 7 | 7 | 156.81 |
| AT5G13480 | 0 | 1 | 1 | 1 | 73.84 |
| AT5G16070 | 0 | 9 | 5 | 8 | 182.67 |
| AT5G18550 | 0 | 1 | 1 | 1 | 40.4 |
| AT5G20890 | 0 | 4 | 3 | 3 | 151.33 |
| AT5G26360 | 0 | 3 | 4 | 3 | 105.5 |
| AT5G28540 | 3 | 13 | 13 | 18 | 209.42 |
| AT5G36230 | 0 | 1 | 1 | 1 | 47.06 |
| AT5G37720 | 0 | 14 | 10 | 11 | 179.44 |
| AT5G42950 | 0 | 11 | 8 | 14 | 204.62 |
| AT5G47010 | 0 | 23 | 12 | 22 | 245.26 |
| AT5G52640 | 5 | 22 | 17 | 19 | 255.25 |
| AT5G54430 | 0 | 3 | 3 | 3 | 111.43 |
| AT5G56010 | 4 | 28 | 26 | 22 | 261.92 |
| AT5G56030 | 4 | 28 | 26 | 22 | 260.7 |
| AT5G59950 | 0 | 5 | 4 | 3 | 89.5 |
| AT5G61140 | 0 | 5 | 4 | 2 | 103.45 |
| AT5G61780 | 0 | 5 | 3 | 4 | 107.34 |
| AT5G62090 | 0 | 2 | 1 | 1 | 72.4 |
| AT5G65250 | 0 | 1 | 2 | 1 | 59.33 |
| AT5G65410 | 0 | 1 | 1 | 1 | 42 |

**Supplementary Table 4 | List of SG-associated§ and 9B-interacting proteins.**

Number of peptides and *P* values are shown.

| Gene id | *GFP* | *9B:GFP* rep1 | *9B:GFP* rep2 | *9B:GFP* rep3 | $-10\log_{10}P$ |
|---|---|---|---|---|---|
| AT1G24510 | 0 | 6 | 8 | 7 | 145.9 |
| AT2G42520 | 0 | 9 | 6 | 8 | 181.9 |
| AT3G09440 | 9 | 37 | 28 | 34 | 263.77 |
| AT3G14100 | 0 | 1 | 1 | 1 | 31.65 |
| AT3G58510 | 0 | 10 | 10 | 11 | 196.36 |
| AT5G20890 | 0 | 4 | 3 | 3 | 151.33 |
| AT3G13460 | 0 | 3 | 2 | 3 | 101.76 |
| AT4G34110 | 2 | 6 | 7 | 9 | 160.5 |
| AT2G23350 | 3 | 10 | 10 | 9 | 184.24 |
| AT1G49760 | 1 | 3 | 4 | 7 | 132.33 |
| AT1G11650 | 0 | 2 | 1 | 2 | 73.33 |
| AT3G19130 | 0 | 2 | 1 | 2 | 60.73 |
| AT4G38740 | 0 | 1 | 1 | 1 | 38.82 |
| AT5G61780 | 0 | 5 | 3 | 4 | 107.34 |
| AT3G13300 | 0 | 1 | 2 | 0 | 62.62 |

§ SG-associated proteins were identified in a previous study by Kosmacz et al., *New Phytologist*, 2019 (doi: 10.1111/nph.15690).
